# The eIF3a subunit drives RNA binding within the eIF3 complex

**DOI:** 10.1101/2024.04.25.581977

**Authors:** Nicholas A. Ide, Riley C. Gentry, Brennan Ashwood, Margaret A. Rudbach, Kyungyoon Yoo, Paloma Kai Velez, Victoria M. Comunale, Erik W. Hartwick, Colin D. Kinz-Thompson, Ruben L. Gonzalez, Colin Echeverría Aitken

**Affiliations:** Department of Biological Sciences, Columbia University, New York, NY, USA; Department of Chemistry, Columbia University, New York, NY, USA; Biochemistry Program, Vassar College, Poughkeepsie, NY, USA; Biology Department, Vassar College, Poughkeepsie, NY, USA; Department of Genetics, Harvard Medical School, Boston, MA, USA; Laboratory of Molecular Biophysics, The Rockefeller University, New York, NY, USA; Renaissance School of Medicine, Stony Brook University, Stony Brook, NY, USA; Biochemistry Krios Electron Microscopy Facility, Department of Biochemistry, University of Colorado Boulder, Boulder, CO, USA; Department of Chemistry, Rutgers University-Newark, Newark, NJ, USA

## Abstract

Eukaryotic initiation factor 3 (eIF3) is a multi-subunit protein complex that binds the 40S pre-initiation complex (PIC) and has an essential role in messenger RNA (mRNA) recruitment, a process encompassing mRNA attachment and loading, scanning, and start-codon recognition. During recruitment, eIF3 is positioned to contact both the 18S ribosomal RNA (rRNA) component of the PIC and the mRNA, but how individual eIF3 subunits contribute to these RNA interactions remains poorly understood. A major obstacle has been the lack of a compositionally defined, biochemically active reconstituted yeast eIF3 complex that enables systematic analysis of individual subunits and defined subcomplexes. Here, we reconstitute Saccharomyces cerevisiae eIF3 from individually purified recombinant subunits and show that it is compositionally stable under our assay conditions and recapitulates core biochemical activities of native eIF3, including PIC binding, stimulation of capped mRNA recruitment, and RNA binding. Using this system, we show that eIF3a provides a core RNA-binding activity within eIF3 and that interactions with other eIF3 subunits modulate this activity in the intact complex and in defined eIF3a-containing subcomplexes. These results establish a reconstituted yeast eIF3 system for dissecting subunit-level biochemical activities and support a model in which subunit context tunes eIF3a-driven RNA binding during initiation.

## INTRODUCTION

Eukaryotic initiation factor 3 (eIF3) is a multi-subunit protein complex essential for forming a productive pre-initiation complex (PIC) on the 40S ribosomal subunit and for supporting messenger RNA (mRNA) translation initiation (Hinnebusch 2006; Aitken and Lorsch 2012; Hinnebusch and Lorsch 2012; Cate 2017). In the conventional cap-dependent initiation pathway, a primary role of eIF3 is to bind the 40S subunit as part of the PIC, where eIF3 facilitates the association of other eIFs and is required for efficient mRNA recruitment (Mitchell et al. 2010; Valasek 2012; Valasek et al. 2017; Wolf et al. 2020; Ide et al. 2025). mRNA recruitment encompasses attachment of the mRNA to the PIC, loading of the mRNA into the mRNA-binding channel of the 40S subunit, scanning of the PIC along the mRNA, and recognition of the start codon. This canonical sequence of events carries an important mechanistic implication. To the extent that eIF3 engages RNA during initiation, the earliest and most proximal RNA contacts are generally expected to involve the 18S ribosomal RNA (rRNA) component of the 40S subunit. Once positioned on the PIC, eIF3 can also engage the incoming mRNA throughout recruitment.

At the same time, eIF3 has been implicated in translational control processes that extend beyond its role as a PIC-associated scaffold (Gomes-Duarte et al. 2018; Wolf et al. 2020). A growing body of work links eIF3 to gene-specific translational outcomes and to activities beyond initiation, including elongation and termination (Beznoskova et al. 2013; Beznoskova et al. 2015; Mohammad et al. 2017; Lin et al. 2020), suggesting that eIF3 can remain associated with, or re-engage, translating ribosomes, at least on select mRNAs. Modular components of eIF3 also occupy positions on the PIC near the mRNA-entry and mRNA-exit channels of the 40S subunit (Erzberger et al. 2014; Llacer et al. 2015; Aitken et al. 2016; Llacer et al. 2021). This positioning is consistent with a model in which the RNA-binding activities of eIF3 contribute both to PIC binding through contacts with rRNA and to mRNA attachment, loading, scanning, and start-codon recognition during recruitment. As predicted by this model, mutations throughout eIF3 alter mRNA-recruitment kinetics under conditions in which PIC binding is saturated, and structural and biochemical studies position eIF3 near mRNA sequences at the exit channel of the PIC (Aitken et al. 2016; Valasek et al. 2017).

More recently, alternative initiation pathways have been proposed in which eIF3 may engage mRNA more directly (Lee, Kranzusch and Cate 2015; Meyer et al. 2015; Hayek et al. 2021; Koubek et al. 2025; Quartey and Goss 2025). In several cap-independent contexts, PIC-free eIF3 or individual eIF3 subunits have been suggested to bind specific mRNAs directly and contribute to their recruitment to the PIC through mRNA-encoded features and/or eIF-bridged interactions (Meyer et al. 2015; Lee et al. 2016; Fujii et al. 2021; Roiuk, Neff and Teleman 2024). Although the generality and mechanistic basis of these pathways remain under active investigation, they suggest that the RNA-binding activities of eIF3 can be used to engage specific mRNAs directly. In addition, the abundance of individual eIF3 subunits varies across cellular conditions, stress, and disease-associated states (Nupponen et al. 1999; Zhang, Pan and Hershey 2007; Herrmannova et al. 2024; Lin et al. 2026), potentially altering the availability or activity of intact eIF3, free subunits, or subcomplexes. Altered expression of individual eIF3 subunits, including subunits conserved in yeast, has been linked to cancer (Zhang, Pan and Hershey 2007; Hershey 2015; Duan et al. 2023) and other disease-associated translational programs (Gomes-Duarte et al. 2018; Jiao et al. 2025). Together, eIF3 engagement of rRNA within the PIC, contact with mRNA during recruitment, and association with ribosomes during later stages of translation suggest that a common RNA-binding capacity could be deployed across multiple modes and stages of translation. Regulated or dysregulated changes in eIF3 subunit abundance or interactions could further alter how this capacity is deployed. Because eIF3 is a modular complex in which individual modules and subunits occupy distinct positions on the 40S subunit (Erzberger et al. 2014; Llacer et al. 2021) and may make different RNA contacts in different contexts, it is essential to understand these contacts at the level of individual subunits and defined subcomplexes. Such an understanding is necessary to develop molecular models explaining how eIF3 engages rRNA within the PIC, supports mRNA recruitment, participates in mRNA-specific alternative initiation pathways, and responds to changes in subunit abundance or interactions.

A major obstacle to investigating the contributions of individual eIF3 subunits to RNA binding is the essential nature of each core subunit, which has precluded genetic deletion of individual components to generate subunit-depleted eIF3 complexes. Previous recombinant eIF3 systems have begun to address this limitation but have undergone different levels of biochemical and functional characterization. A mammalian eIF3 complex assembled from recombinant subunits expressed in *Escherichia coli* was shown to bind the hepatitis C virus internal ribosome entry site (IRES) and other PIC components and to assemble into IRES-mediated translation complexes in extract (Sun et al. 2011). Previous recombinant yeast eIF3 preparations were used for structural studies or were shown to rescue translation in a heat-inactivated extract containing a temperature-sensitive eIF3b mutant, but they were not compared with native eIF3 across PIC binding, mRNA recruitment, and direct RNA binding (Khoshnevis et al. 2012; Llacer et al. 2015; Zeman et al. 2019). Establishing a compositionally defined yeast eIF3 complex that recapitulates these core biochemical activities would offer significant advantages. Yeast eIF3 consists of a conserved five-subunit complex that captures fundamental eIF3 functions in a simpler, more experimentally tractable architecture corresponding to the conserved core scaffold within the larger mammalian complex (Valasek et al. 2017; Ide et al. 2025). A fully functional recombinant yeast eIF3 can therefore provide a reference system for defining the activities of the conserved core. In particular, such a system enables investigation of how individual core subunits engage rRNA within the PIC and mRNA during recruitment, providing a basis for interpreting the contributions of additional mammalian subunits to these processes.

Here, we report the reconstitution of *Saccharomyces cerevisiae* eIF3 from individually purified recombinant subunits and demonstrate that reconstituted eIF3 is compositionally stable under our assay conditions and displays PIC-binding, mRNA-recruitment, and RNA-binding activities comparable to those of native *S. cerevisiae* eIF3. Using this compositionally defined and functionally validated system, we investigate how eIF3 binds RNA and determine the contributions of individual eIF3 subunits and defined subcomplexes to this activity. We show that RNA binding by eIF3 is driven predominantly by eIF3a and that interactions between eIF3a and other eIF3 subunits modulate this activity. Collectively, our results support a mechanistic framework in which subunit-subunit interactions within eIF3 tune a core, eIF3a-driven RNA-binding activity. This framework provides a foundation for understanding how eIF3 supports mRNA recruitment and how regulated changes in eIF3 subunit abundance, interactions, or assembly state could influence eIF3 function across translation.

## RESULTS

### Reconstitution of recombinant yeast eIF3

Because each core eIF3 subunit is essential for cell survival, mechanistic interrogation of the activities of individual subunits *in vitro* has largely been limited to viable mutations. For example, one study used a suite of mutations across eIF3 subunits to identify defects in eIF3 activity *in vitro* (Aitken et al. 2016). Other studies focused on the activities of additional eIFs or on translation initiation more broadly have used wild-type eIF3 purified directly from *S. cerevisiae* (Mitchell et al. 2010; Yourik et al. 2017; Liu et al. 2019). Notably, the biochemical activity of natively purified eIF3 can vary substantially among preparations (Liu et al. 2019). Only three studies have attempted to generate recombinant *S. cerevisiae* eIF3 using subunits expressed in *Escherichia coli*. One study co-expressed all five eIF3 subunits for structural studies but reported subunit degradation (Llacer et al. 2015). The other two studies individually expressed and purified the subunits and demonstrated that they could rescue a heat-inactivated cell extract containing a temperature-sensitive, loss-of-function eIF3b mutant (Khoshnevis et al. 2012; Zeman et al. 2019). None of these recombinant systems was shown, in the absence of native eIF3 subunits, to recapitulate core biochemical activities of eIF3, such as PIC binding or stimulation of mRNA recruitment. Together with the variation observed among native eIF3 preparations (Mitchell et al. 2010; Aitken et al. 2016; Yourik et al. 2017), this limitation complicates direct comparison of eIF3 activities and underscores the challenges of studying individual eIF3 subunits and defined subcomplexes.

To establish an eIF3 complex suitable for studying individual subunits and defined subcomplexes, we reconstituted eIF3 from individually purified recombinant subunits (Figure 1A). We purified eIF3a, eIF3b, and eIF3c from *E. coli*. We also purified an eIF3i-eIF3g fusion from *E. coli* and separated the eIF3i and eIF3g subunits by proteolytic cleavage, yielding a co-purified eIF3ig subcomplex according to a previously described method for preparing reconstituted *S. cerevisiae* eIF3 (Zeman et al. 2019). Combining these purified subunits in stoichiometric amounts yielded reconstituted *S. cerevisiae* eIF3 (eIF3_R_), which eluted from a size-exclusion chromatography column as a single peak with a profile similar to that of native *S. cerevisiae* eIF3 (eIF3_N_) (Figure 1B).

**Figure 1.**
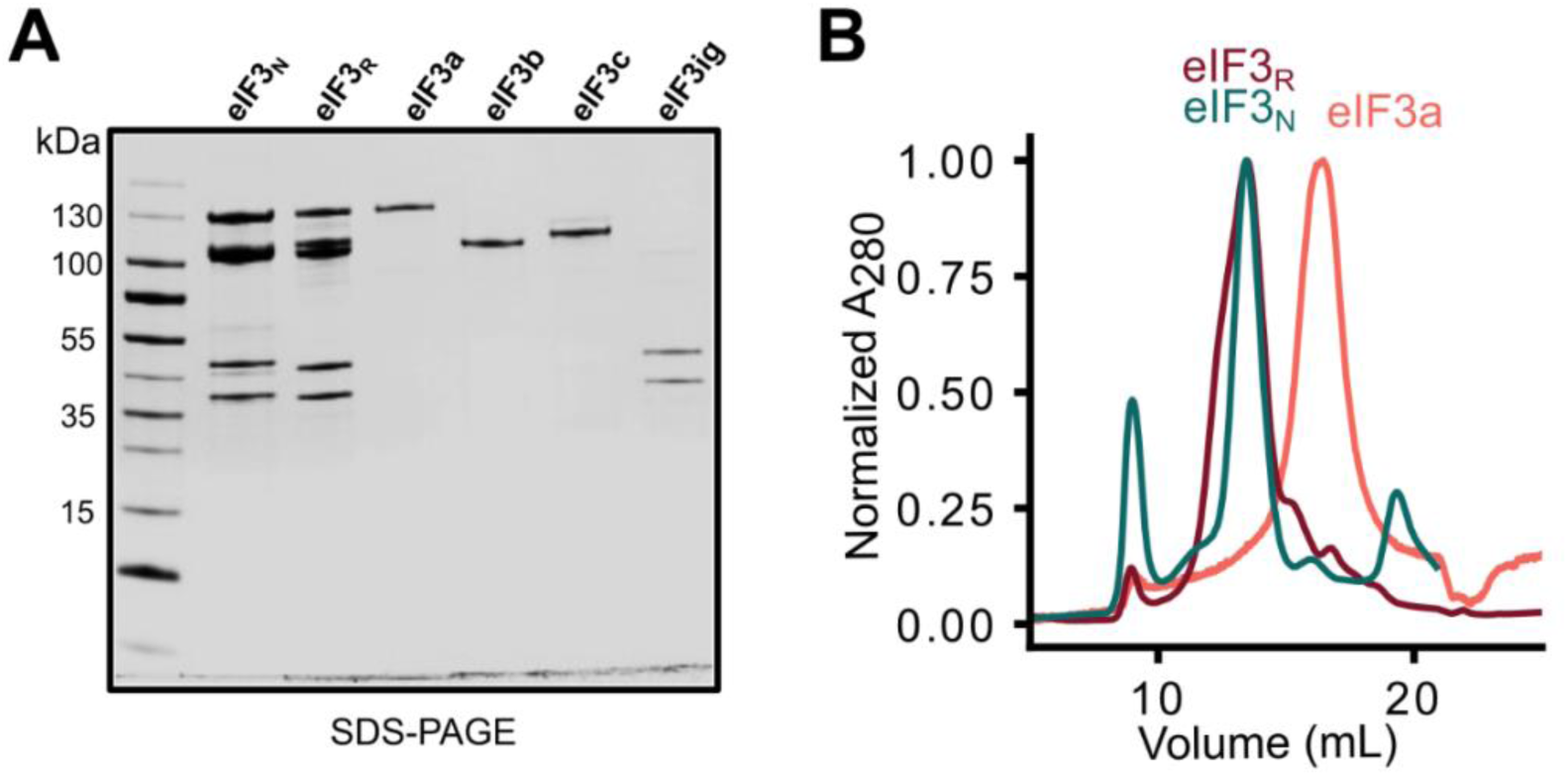
Preparation of native and reconstituted yeast elF3. **(A)** SDS-PAGE analysis of elF3_N_, eIFS_r_, and the recombinant elF_3_ subunits or subcomplexes used for reconstitution. **(B)** Absorbance size-exclusion chromatograms comparing eIF3_n_, eIF3_r_, and isolated elF3*a*. Absorbance at 280 nm was normalized to the maximum signal for each chromatogram. eIF3r elutes predominantly as a single peak with a profile similar to that of eIF3_n_, consistent with assembly of the recombinant subunits into an intact elF_3_ complex.

We next assessed the compositional stability of eIF3_R_ in solution using fluorescence-tracked size-exclusion chromatography assays that monitor exchange of a fluorophore-labeled eIF3b subunit. We labeled a single-cysteine variant of eIF3b with Janelia Fluor 549 ((JF549)eIF3b; Figure S1A), assembled a (JF549)eIF3b-containing eIF3_R_ complex, and analyzed its elution profile over a concentration range of 30 to 300 nM (Figure S1B). Across this range, the elution profile of the (JF549)eIF3b-containing eIF3_R_ complex was unchanged and remained dominated by a single peak corresponding to the intact complex (Figure S1B). We then monitored incorporation of (JF549)eIF3b, either alone or in complex with eIF3ig, into preformed, unlabeled eIF3_R_ (Figures S2A and S2B). Exchange of (JF549)eIF3b into eIF3_R_ occurred slowly, with maximal incorporation occurring over hours (Figures S2A and S2C). The (JF549)eIF3big subcomplex, consisting of (JF549)eIF3b in complex with eIF3ig, incorporated more rapidly and to a greater extent than (JF549)eIF3b alone but still required more than an hour to achieve maximal exchange (Figures S2B and S2C). Taken together, these results indicate that eIF3_R_ is compositionally stable under our experimental conditions and that exchange of individual subunits and subcomplexes occurs slowly, arguing against substantial exchange on the timescale of the biochemical assays described below. These findings therefore support the use of eIF3 _R_ as a compositionally defined reference complex for comparison with individual subunits and defined subcomplexes.

We also collected conventional mass photometry measurements at low nanomolar concentrations. Because conventional mass photometry detects molecules through surface-landing events, interpreting detected event distributions as quantitative solution-phase subpopulation distributions depends on assumptions regarding diffusion, surface landing, analyte concentration, surface chemistry, and detection that have been discussed extensively in the mass photometry methodology literature (Sonn-Segev et al. 2020; Marciano et al. 2022; Asor et al. 2025; Kratochvil et al. 2026). Under our experimental conditions, the resulting surface-associated event distributions could not be reconciled with the solution-phase stability and exchange behavior observed by fluorescence-tracked size-exclusion chromatography. We therefore present these data in the Supplementary Materials and conservatively treat them as non-diagnostic for the solution-phase compositional stability of eIF3 under our conditions (Figure S3).

### eIF3_R_ recapitulates the biochemical activities of eIF3_N_

Having established that eIF3_R_ exhibits a size-exclusion chromatography profile similar to that of eIF3_N_ and is compositionally stable under our assay conditions, we compared eIF3_N_ and eIF3_R_ across three established biochemical activities of eIF3: (*i*) PIC binding, (*ii*) stimulation of capped mRNA recruitment to the PIC, and (*iii*) direct RNA binding. First, to compare PIC binding by eIF3_N_ and eIF3_R_, we employed a previously described *S. cerevisiae* reconstituted translation-initiation system (Acker et al. 2007; Aitken et al. 2016; Liu et al. 2019; Gentry et al. 2025) to assemble PICs (Methods). To visualize these PICs, we used a previously described uncapped and unstructured 50-nucleotide model mRNA containing an AUG start codon at nucleotides 23-25 and a fluorescein label at the 3′ end (Aitken et al. 2016). Briefly, we combined this mRNA and 40S subunits with saturating concentrations of eIF1, eIF1A, and a ternary complex composed of eIF2, guanosine 5′-[β,γ-imido]triphosphate (GDPNP), and initiator tRNA (eIF2(GDPNP)tRNA_i_) to form PICs under conditions previously shown to support start-codon-dependent PIC formation in the absence of eIF4 factors. We then incubated these PICs with varying concentrations of either eIF3_N_ or eIF3_R_ and used native polyacrylamide gel electrophoresis to separate free mRNA, eIF3-free PICs, and eIF3-bound PICs (Figure 2A). Consistent with a previous study reporting a mean apparent equilibrium dissociation constant (*K*_d_^app^) of ≤ 38 nM for eIF3_N_ binding to the PIC (Aitken et al. 2016), we measured mean *K*_d_^app^ values of 23 nM for eIF3_N_ and 13 nM for eIF3_R_ (Figure 2A). Thus, eIF3_R_ binds the PIC with an apparent affinity comparable to that of eIF3_N_.

**Figure 2.**
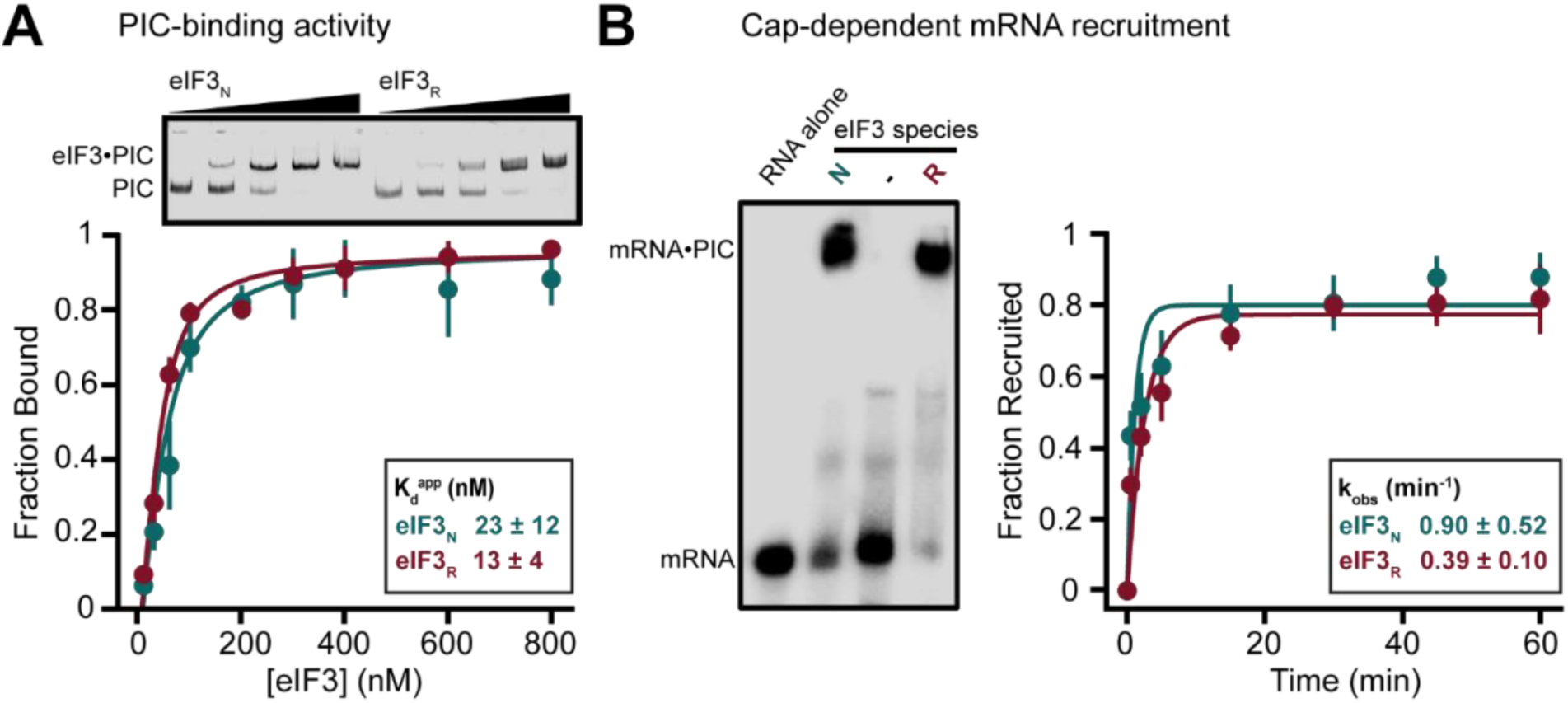
eIF3_R_ recapitulates the PIC-binding and mRNA-recruitment activities of eIF3_N_. **(A)** Electrophoretic mobility-shift assay (EMSA) comparing binding of eIF3_N_ and eIF3_R_ to PICs assembled on a fluorescein-labeled model mRNA. A representative gel is shown above the quantified fraction of eIF_3_-bound PICs. Points represent the mean of three independent experiments, error bars represent the standard deviation, and solid curves show fits to the quadratic binding equation (Methods). Each K_d_^app^ is indicated as a mean and a standard deviation. **(B)** EMSA comparing the abilities of eIF3_N_ and eIF3_R_ to stimulate recruitment of capped ^32^P-labeled rpl41a mRNA into PICs. Because formation of the gel-shifted complex depends on start-codon recognition, this assay reports on completion of the mRNA-recruitment process. A representative gel is shown alongside the quantified fraction of recruited mRNA. Points represent the mean of three independent experiments, error bars represent the standard deviation, and solid curves show fits to a single­exponential equation (Methods). Each k_obs_ is indicated as a mean and a standard deviation.

Next, we compared the abilities of eIF3_N_ and eIF3_R_ to stimulate *in vitro* recruitment of capped mRNA to PICs using a previously established assay (Mitchell et al. 2010; Aitken et al. 2016; Yourik et al. 2017). This assay monitors recruitment of a capped mRNA encoding the *S. cerevisiae* rpl41a gene to PICs as a function of time. The rpl41a mRNA has been used as a representative natural mRNA in numerous *in vitro* and *in vivo* biochemical studies (Nielsen et al. 2004; Jivotovskaya et al. 2006; Chiu et al. 2010; Mitchell et al. 2010; Khoshnevis et al. 2014; Aitken et al. 2016). Briefly, we prepared reaction mixtures containing 40S subunits; saturating concentrations of eIF1, eIF1A, and eIF2(GDPNP)tRNA_i_; the specified concentration of either eIF3_N_ or eIF3_R_; saturating concentrations of the eIF4 factors eIF4G, eIF4E, eIF4A, and eIF4B; and ATP (Methods). We initiated each reaction by adding a substoichiometric concentration of rpl41a mRNA containing a ^32^P-labeled cap. After incubation for the indicated times, we used native polyacrylamide gel electrophoresis to separate free mRNA from mRNA recruited to PICs. At saturating concentrations of eIF3, eIF3_N_ and eIF3_R_ both supported rapid mRNA recruitment, with observed rate constants of 0.90 ± 0.52 min^−1^ and 0.39 ± 0.10 min^−1^, respectively, and comparable recruitment endpoints over the 60-minute time course (Figure 2B). Across eIF3 concentrations ranging from near-equimolar to saturating relative to the PIC, both eIF3_N_ and eIF3_R_ supported recruitment of > 70% of the mRNA (Figure S4). Collectively, these results demonstrate that eIF3_R_ recapitulates the established ability of eIF3_N_ to stimulate *in vitro* recruitment of capped mRNA to PICs.

Lastly, we used the same uncapped, 50-nucleotide, 3′-fluorescein-labeled model RNA employed in the PIC-binding assays to compare direct RNA binding by eIF3_N_ and eIF3_R_. We measured changes in RNA fluorescence anisotropy upon titration with either eIF3_N_ or eIF3_R_ (Figure 3A). Fitting the resulting data to a model in which a single eIF3 species binds a single RNA site yielded mean *K* ^app^ values of 5 nM for eIF3_N_ and 47 nM for eIF3_R_ (Figure 3A). The eIF3_R_ value agrees closely with the mean *K*_d_^app^ of 35 nM previously reported for binding of eIF3 to RNA (Mitchell et al. 2010). Although subsequent experiments show that RNA binding by intact eIF3 does not conform fully to a two-component, single-site model (see below), these apparent values establish that both eIF3_N_ and eIF3_R_ bind RNA with high affinity. Thus, eIF3_R_ recapitulates the direct RNA-binding activity of eIF3_N_.

**Figure 3.**
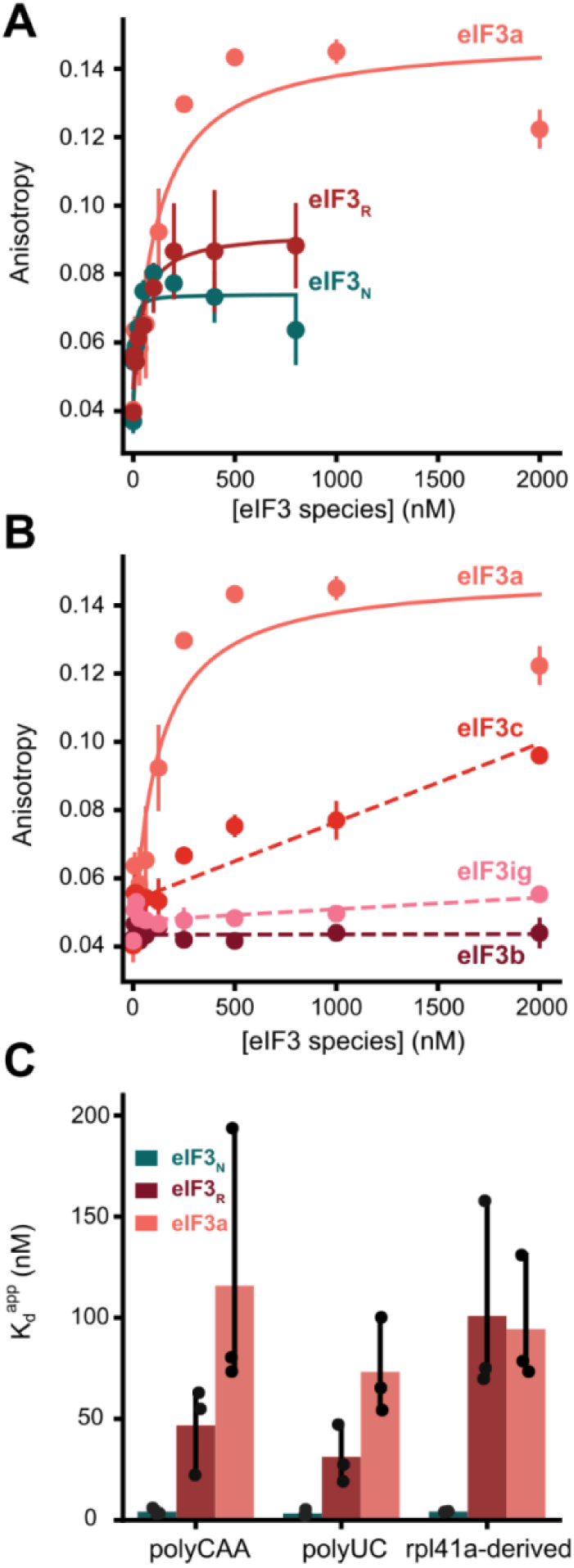
elF3a is the only isolated elF3 subunit tested that binds RNA with high affinity. **(A)** Fluorescence-anisotropy measurements comparing binding of elF3_N_, eIF3_r_, and isolated elF3a to a fluorescein-labeled polyCAA model RNA. Points represent mean anisotropy values from three independent titrations, and bars represent the standard deviation. Solid curves indicate fits to the quadratic binding equation (Methods). **(B)** Fluorescence-anisotropy measurements comparing the binding of isolated elF3a, elF3b, elF3c, and the co-purified elF3ig subcomplex to the same model RNA. Data are represented as in Panel A. The elF3a dataset is repeated from Panel A for comparison. elF3c bound RNA weakly and did not reach saturation over the concentration range tested, whereas no appreciable binding by elF3b or elF3ig was detected. Dashed lines are included only as visual guides and do not represent binding-model fits. **(C)** K_d_^app^s measured for binding of 6IF3_n_, oIF3_r_, and elF3a to polyCAA, polyUC, and rpl41 a-derived model RNAs. Bars represent the mean K_d_^app^ values obtained from three independent titrations, individual values are shown as points, and error bars represent the standard deviation. The similar apparent affinities measured across the three RNAs indicate no strong sequence preference among the sequences tested.

Although eIF3_R_ recapitulates the core biochemical activities of eIF3_N_, quantitative differences between the two preparations are apparent across the assays. Such differences are consistent with previous observations that biochemical activity can vary among native eIF3 preparations and could reflect factors that differentially influence PIC binding, mRNA recruitment, and ensemble RNA-binding measurements. For example, eIF3_N_ preparations may contain co-purifying proteins and post-translational modifications that are absent from eIF3_R_ and may exhibit subtle variation in subunit stoichiometry relative to the more stoichiometrically uniform eIF3 _R_. Nonetheless, these assays establish that eIF3_R_ recapitulates PIC binding, stimulation of capped mRNA recruitment, and direct RNA binding by eIF3_N_, thereby providing a compositionally defined system for dissecting the contributions of individual subunits and defined subcomplexes to RNA binding.

### eIF3a is the only free eIF3 subunit that binds RNA with high affinity

To identify which subunits might contribute to RNA binding by intact eIF3, we leveraged our reconstituted system to test whether and to what extent individual eIF3 subunits interact directly with RNA in isolation. We adapted the fluorescence anisotropy-based RNA-binding assay described above to measure changes in RNA fluorescence anisotropy upon titration with individual eIF3 subunits (Figures 3B, S5, S6B, and Table S1). Because eIF3i and eIF3g co-purify as the eIF3ig subcomplex, we tested these two subunits together. Despite the presence of RNA-recognition motifs in both eIF3b and eIF3g (Hanachi, Hershey and Vornlocher 1999; ElAntak et al. 2007; Khoshnevis, Neumann and Ficner 2010), neither eIF3b nor eIF3ig exhibited appreciable RNA binding, even at the highest protein concentrations tested (2 µM) (Figures 3B, S5, and Table S1). eIF3c exhibited only modest affinity for RNA, with a *K*_d_ ^app^ > 1000 nM (Figures S5B and S6B and Table S1). In contrast, eIF3a bound RNA with high affinity, yielding a mean *K*_d_^app^ of 116 nM and identifying eIF3a as the only individual eIF3 subunit tested that binds RNA with high affinity (Figures 3B and S5 and Table S1). Thus, eIF3a provides a core RNA-binding activity that likely contributes to the activity observed for eIF3_N_ and eIF3_R_, which yielded mean *K*_d_^app^ values of 5 and 45 nM, respectively. Although eIF3b, eIF3c, and eIF3ig did not display high-affinity RNA binding under our experimental conditions, the greater apparent RNA-binding affinities of eIF3_N_ and eIF3_R_ relative to eIF3a alone suggest that the additional subunits collectively modulate eIF3a -driven RNA binding.

Among the RNA lengths tested, high-affinity binding by eIF3_N_, eIF3_R_, and eIF3a was observed only for RNAs of at least 45 nucleotides (Table S1). To investigate the influence of RNA sequence, we tested a model RNA containing a distinct repetitive sequence (Figures 3C, S5A, and S5B and Table S1) and a second model RNA containing the first 51 nucleotides of the rpl41a mRNA sequence (Figures 3C, S5C, and S5D and Table S1). Both additional RNAs yielded RNA-binding behavior comparable to that observed with the initial model RNA (Figures 3 and S5 and Table S1). Thus, the eIF3 species tested showed no strong sequence preference among these three RNAs in the absence of additional interacting components.

Because eIF3c displayed weak independent RNA-binding activity and the C-terminal domain of eIF3c forms a heterodimer with the N-terminal domain of eIF3a that stabilizes mRNA binding to the PIC (Khoshnevis et al. 2014; Aitken et al. 2016), we asked whether eIF3c modulates RNA binding by eIF3a. Inclusion of an equimolar concentration of eIF3c with eIF3a yielded a binding curve more similar to that of eIF3_R_ than to that of isolated eIF3a, indicating that association of eIF3c with eIF3a, presumably within an eIF3ac complex, alters eIF3a-driven RNA binding (Figure S6A). Whether RNA binding by eIF3ac reflects the combined RNA-binding activities of eIF3a and eIF3c or a change in eIF3a-driven RNA binding caused by formation of the eIF3ac complex remains unclear. Nevertheless, these results demonstrate that interaction with eIF3c modulates eIF3a-driven RNA binding within a defined subcomplex and support a model in which subunit context modulates this activity within intact eIF3.

### Subunit context modulates eIF3a-driven RNA binding

Given that isolated eIF3a and the eIF3ac subcomplex both bind RNA, we next sought to determine how eIF3a-driven RNA binding is expressed within intact eIF3 and whether defined eIF3a-containing subcomplexes are also competent to engage RNA. If RNA binding by eIF3 involves multiple binding modes, multiple binding-competent species, or both, then a two-component, single-site binding model may be inappropriate for analyzing ensemble titrations. Such models have nonetheless been widely used as simplifying assumptions in previous biochemical studies of eIF3. By comparison, RNA binding by isolated eIF3a is expected to approximate a bimolecular interaction more closely than RNA binding by eIF3R or eIF3N. We therefore tested whether RNA binding by intact eIF3 diverges from the behavior expected for a single binding species interacting with a single RNA site.

We first performed fluorescence-anisotropy experiments in which the total RNA concentration was varied across independent titrations. Under a two-component, single-site binding model, increasing the RNA concentration is expected to shift the binding curve toward higher protein concentrations and, at RNA concentrations near or above *K*_d_, produce a non-hyperbolic titration curve (Jarmoskaite et al. 2020). Because we can define the subunit stoichiometry of eIF3_R_ but not eIF3_N_, we compared eIF3a and eIF3_R_ in these experiments. As expected, titrations with eIF3a behaved in a manner consistent with a two-component, single-site binding model, with higher RNA concentrations requiring proportionally higher eIF3a concentrations to saturate binding (Figure 4, left). Equivalent titrations with eIF3_R_ did not exhibit this behavior (Figure 4, right). Instead, the anisotropy reached a maximum at approximately 500 nM eIF3_R_, largely independently of the RNA concentration, and then decreased at higher eIF3 _R_ concentrations. We observed a similar decrease in anisotropy at high concentrations of eIF3_N_ (Figure 3A). Together with the fluorescence-tracked size-exclusion chromatography experiments showing that eIF3_R_ is compositionally stable on the timescale of these assays, these observations suggest that RNA binding by intact eIF3 does not behave as a simple two-component, single-site interaction. Instead, other eIF3 subunits may modulate eIF3a-driven RNA binding by contributing additional RNA contacts, altering the accessibility or geometry of the eIF3a RNA-binding surface, or promoting transitions among distinct RNA-bound states. Low-abundance eIF3a-containing subcomplexes or free eIF3a in the preparation could also contribute to the ensemble signal, although our compositional-stability experiments argue against substantial exchange or accumulation of these species during the assay. Thus, the ensemble anisotropy measurements most likely reflect multiple RNA-binding modes within intact eIF3, while possible contributions from low-abundance eIF3a-containing species cannot be excluded.

**Figure 4.**
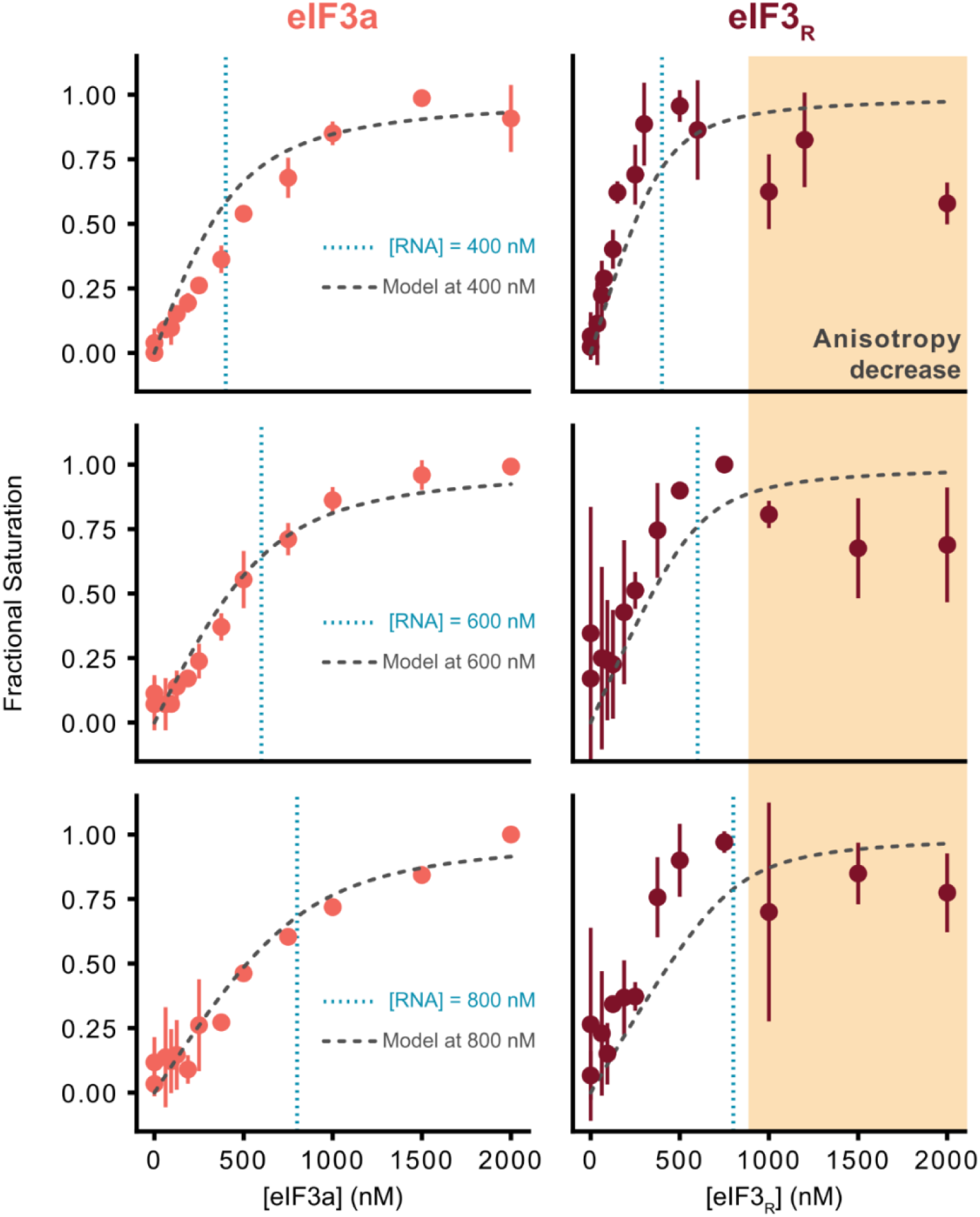
RNA binding by elF3R diverges from a two-component, single-site binding model. Fluorescence-anisotropy binding measurements comparing RNA binding by isolated elF3a and eIF3_r_ at total polyCAA RNA concentrations of 400, 600, or 800 nm, as indicated. Anisotropy values were converted to fractional saturation. Gray dashed curves show the behavior predicted by a two-component, single-site binding model using the K_d_^app^ values measured in *Figure 3*, and blue dotted lines mark the corresponding total RNA concentrations. Points represent the mean of three independent titrations, and error bars represent the standard deviation. Increasing RNA concentration shifted the elF3a titrations to proportionally higher protein concentrations, consistent with the model. In contrast, the elF3R titrations deviated from the predicted behavior and showed decreased anisotropy at high eIFS_r_ concentrations. Together with the compositional stability of elF3_R_ on the timescale of these measurements, these results indicate that RNA binding by intact elF3_R_ is not adequately described as a single two-component, single-site interaction and is consistent with the presence of multiple RNA-binding modes.

We next asked whether altered expression of individual eIF3 subunits produces cellular phenotypes. Under normal cellular conditions, the stoichiometry of eIF3 subunits in *S. cerevisiae* is expected to be nearly equal (von der Haar and McCarthy 2002; Ho, Baryshnikova and Brown 2018), such that intact five-subunit eIF3 is expected to be the predominant species. To perturb this stoichiometry, we cloned each eIF3 subunit into overexpression plasmids containing either the constitutive high-expression TEF1 promoter or the lower-expression RNR2 promoter, introduced these plasmids into an otherwise wild-type *S. cerevisiae* strain, and monitored growth using a spotting assay (Figure S7). Under the TEF1 promoter, expression of every subunit other than eIF3g slowed growth, and transformants expressing eIF3a could not be propagated (Figure S7A). Under the RNR2 promoter, the growth effects were substantially reduced (Figure S7B). These results demonstrate that altered expression of individual eIF3 subunits can perturb cellular fitness. Although the strong phenotype associated with eIF3a expression is notable given the RNA-binding activity of this subunit, the mechanisms underlying these phenotypes are likely multifactorial and may differ among subunits. The experiments therefore do not distinguish among effects on intact eIF3 assembly or abundance, accumulation of free subunits or subcomplexes, altered RNA binding, or other eIF3-dependent processes.

Although our anisotropy experiments suggest that RNA binding by intact eIF3 involves multiple RNA-binding modes, they cannot establish whether the ensemble signal arises exclusively from intact eIF3 or includes contributions from low-abundance eIF3a-containing species. To characterize the apparent-mass classes associated with RNA-dependent binding events and to determine whether defined eIF3a-containing subcomplexes can engage RNA, we adapted a tethered-RNA mass photometry approach. In this assay, we derivatize a borosilicate glass surface with a mixture of polyethylene glycol (PEG) and biotinylated PEG, similar to flow cells used for single-molecule fluorescence microscopy (Figure 5A) (Roy, Hohng and Ha 2008; Gentry et al. 2025; Gentry et al. 2026). This derivatization enables sparse tethering of a 3′-biotinylated version of the model RNA used in the PIC-binding and ensemble RNA-binding experiments through a biotin-streptavidin-biotin bridge. The tethered-RNA format is used here to identify apparent-mass classes associated with repeated binding events at individual RNA-containing locations and is conceptually distinct from the conventional mass photometry measurements described above. The PEG-derivatized surface differs chemically from the bare borosilicate glass used in conventional mass photometry and may therefore alter the nature and frequency of nonspecific surface-associated events. Nonetheless, the tethered-RNA assay remains surface-dependent and we do not use it to infer solution-phase eIF3 composition, stability, or subunit populations. Instead, RNase sensitivity and enrichment for repeated association-dissociation events provide complementary operational criteria for identifying RNA-dependent events. A practical limitation is that the detection laser heats the derivatized surface and likely perturbs the PEG layer, requiring acquisition of a larger number of shorter movies than are used with bare borosilicate coverslips (Figure S8A and Methods).

**Figure 5.**
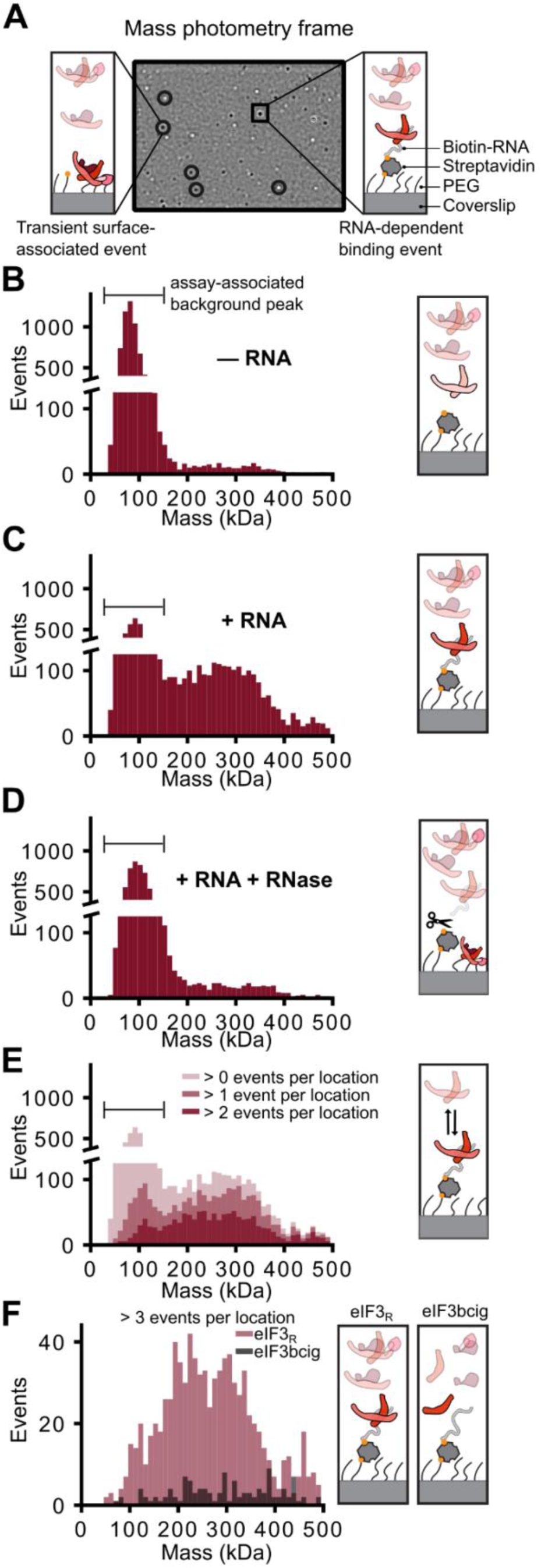
Tethered-RNA mass photometry identifies elF3a-dependent RNA-binding events. **(A)** Representative mass-photometry frame and schematic illustrating transient surface-associated events and RNA-dependent binding events on PEG-derivatized coverslips bearing sparsely tethered biotinylated RNA. **(B-D)** Apparent-mass distributions obtained after delivery of200 nM eIF3_r_ to PEG-derivatized coverslips **(B)** in the absence of tethered RNA, **(C)** in the presence of tethered RNA, or **(D)** after RNase treatment of tethered RNA. A prominent apparent-mass population centered near 100 kDa was observed in the absence of tethered RNA and persisted after RNase treatment and was therefore treated as assay-associated background. Addition of tethered RNA produced a broader, higher-mass distribution that was largely eliminated by RNase treatment. **(E)** Apparent-mass distributions for events detected in the presence of tethered RNA after progressively restricting the analysis to locations exhibiting more than zero, more than one, or more than two events. Increasingly stringent filtering depleted the assay-associated background population while retaining the broader RNA-dependent distribution, consistent with repeated association and dissociation at tethered RNA molecules. **(F)** Apparent-mass distributions for location exhibiting more than three events after delivery of eIF3_r_ or an elF3bcig mixture containing elF3b, elF3c, and elF3ig but lacking elF3a. The marked reduction in repeated events in the absence of elF3a indicates that elF3a is a major determinant of the robust RNA-dependent events detected in this assay. These apparent-mass distributions were not used to infer solution-phase elF3 composition or binding affinities.

We began by comparing events detected after delivery of 200 nM eIF3_R_ to PEG-derivatized wells in the absence or presence of tethered RNA. Under both conditions, the apparent-mass distribution contained a pronounced peak centered at approximately 150 kDa (Figures 5B and 5C). Because this peak was also present in buffer-only controls and did not correspond to the expected mass of an eIF3 subunit, defined subcomplex, or intact eIF3, we treated it as an assay-associated background feature of the PEG-derivatized format. In the presence of tethered RNA, we additionally observed a broader population centered at an apparent mass of approximately 250 kDa that we hypothesized reflected RNA-dependent events (Figure 5C). We tested this hypothesis in two ways. First, we reasoned that RNA-dependent events should be sensitive to ribonuclease (RNase) treatment. Consistent with this prediction, RNase treatment largely eliminated the broader population and produced a distribution closely resembling that observed in the absence of tethered RNA (Figures 5B and 5D). Second, we reasoned that locations containing tethered RNA should exhibit repeated association and dissociation events, whereas nonspecific surface-associated events should more frequently occur as isolated events. We therefore filtered the data to retain locations exhibiting multiple events. Consistent with this prediction, increasingly stringent multi-event filtering preferentially depleted the approximately 150-kDa background population while retaining the broader apparent-mass distribution observed in the presence of tethered RNA (Figure 5E). The RNase-sensitive and multi-event analyses therefore yielded similar apparent-mass distributions, supporting the conclusion that the broader population represents RNA-dependent events. This distribution overlapped the expected mass ranges of both eIF3 subcomplexes and intact eIF3, consistent with heterogeneous RNA-dependent events but insufficient to assign the underlying molecular species uniquely.

We next tested whether robust, repeated RNA-binding events depend on eIF3a. Because ensemble anisotropy experiments showed that eIF3a is the only free subunit that binds RNA with high affinity (Figure 3), we predicted that removal of eIF3a would substantially reduce repeated events in the tethered-RNA assay. We therefore performed tethered-RNA mass photometry experiments using a mixture containing eIF3b, eIF3c, eIF3i, and eIF3g at 200 nM each but lacking eIF3a (eIF3bcig). Comparison of the RNA-dependent event distributions obtained with eIF3_R_ and eIF3bcig using the same multi-event filtering strategy showed that robust, repeated RNA-binding events depended strongly on the presence of eIF3a (Figure 5F). Because the multi-event filter is stringent, this analysis is expected to undercount weak, rare, or transient binding events, including those expected from the weak RNA-binding activity of eIF3c (Figure 3). These results support the conclusion that eIF3a is a major determinant of the robust RNA-dependent events detected in this assay. They do not, however, establish the relative contributions of intact eIF3 and low-abundance subcomplexes within an eIF3_R_ sample.

To determine whether defined eIF3a-containing subcomplexes are competent to engage RNA, we performed additional tethered-RNA experiments using 200 nM eIF3a, eIF3ac, eIF3ab, eIF3abc, or eIF3abig. Each sample produced repeated RNA-binding events with apparent-mass distributions that overlapped the expected mass range of the corresponding protein or subcomplex (Figures S8B–S8F). These results demonstrate that isolated eIF3a and multiple defined eIF3a-containing subcomplexes can engage tethered RNA under these conditions. Because the current mass photometry format does not support robust determination of binding affinities from event counts alone, we do not infer a thermodynamic rank ordering among these species. Collectively, the ensemble anisotropy and tethered-RNA measurements support a model in which eIF3a provides a core RNA-binding activity and interactions with other eIF3 subunits modulate how this activity is expressed within intact eIF3 and across defined eIF3a-containing subcomplexes.

## DISCUSSION

Here, we establish a recombinant *S. cerevisiae* eIF3 complex assembled from individually purified subunits and show that it recapitulates core biochemical activities of native eIF3. Specifically, eIF3_R_ binds the PIC, stimulates *in vitro* recruitment of capped mRNA to PICs, and binds RNA with activities comparable to those of eIF3_N_. In addition, fluorescence-tracked size-exclusion chromatography and subunit-exchange experiments show that eIF3_R_ is compositionally stable under our assay conditions, supporting its use as a compositionally defined reference complex for dissecting the contributions of individual eIF3 subunits and defined subcomplexes. This system provides a framework for investigating how the conserved core subunits of eIF3 contribute to PIC binding, mRNA recruitment, and other modes of RNA engagement during translation.

Using this system, we find that eIF3a provides a core RNA-binding activity within eIF3 (Figure 6). Among the individual eIF3 subunits tested, eIF3a is the only free subunit that binds RNA with high affinity (Figure 6A), despite the presence of canonical RNA-recognition motifs in other eIF3 subunits (Hanachi, Hershey and Vornlocher 1999; ElAntak et al. 2007; Khoshnevis, Neumann and Ficner 2010). The apparent RNA-binding affinity of isolated eIF3a is lower than that of intact eIF3, indicating that interactions with other eIF3 subunits modulate eIF3a-driven RNA binding. Consistent with this conclusion, inclusion of eIF3c alters the RNA-binding behavior of eIF3a such that the eIF3ac subcomplex behaves more similarly to intact eIF3 than to isolated eIF3a (Figures S6A and 6B). Ensemble fluorescence-anisotropy measurements further show that RNA binding by eIF3_R_ does not behave as a simple two-component, single-site interaction. In contrast to isolated eIF3a, which behaves in a manner consistent with such a model, eIF3_R_ exhibits RNA-concentration-dependent behavior that deviates from the expectations for a single binding species, and both eIF3_R_ and eIF3_N_ exhibit decreases in anisotropy at high eIF3 concentrations. Given that eIF3_R_ is compositionally stable on the timescale of these assays, the most likely interpretation is that RNA binding by intact eIF3 involves multiple RNA-binding modes (Figure 6B). These modes could arise from additional RNA contacts contributed by other eIF3 subunits, changes in the accessibility or geometry of the eIF3a RNA-binding surface, transitions among distinct RNA-bound states of the complex, or some combination of these mechanisms. Thus, rather than reporting a single affinity for a single RNA-bound state, ensemble RNA-binding measurements of intact eIF3 likely reflect a distribution of RNA-bound conformations or engagement modes. Possible contributions from low-abundance eIF3a-containing subcomplexes or free eIF3a cannot be excluded, but our compositional-stability experiments argue against substantial exchange or accumulation of these species during our assays.

**Figure 6.**
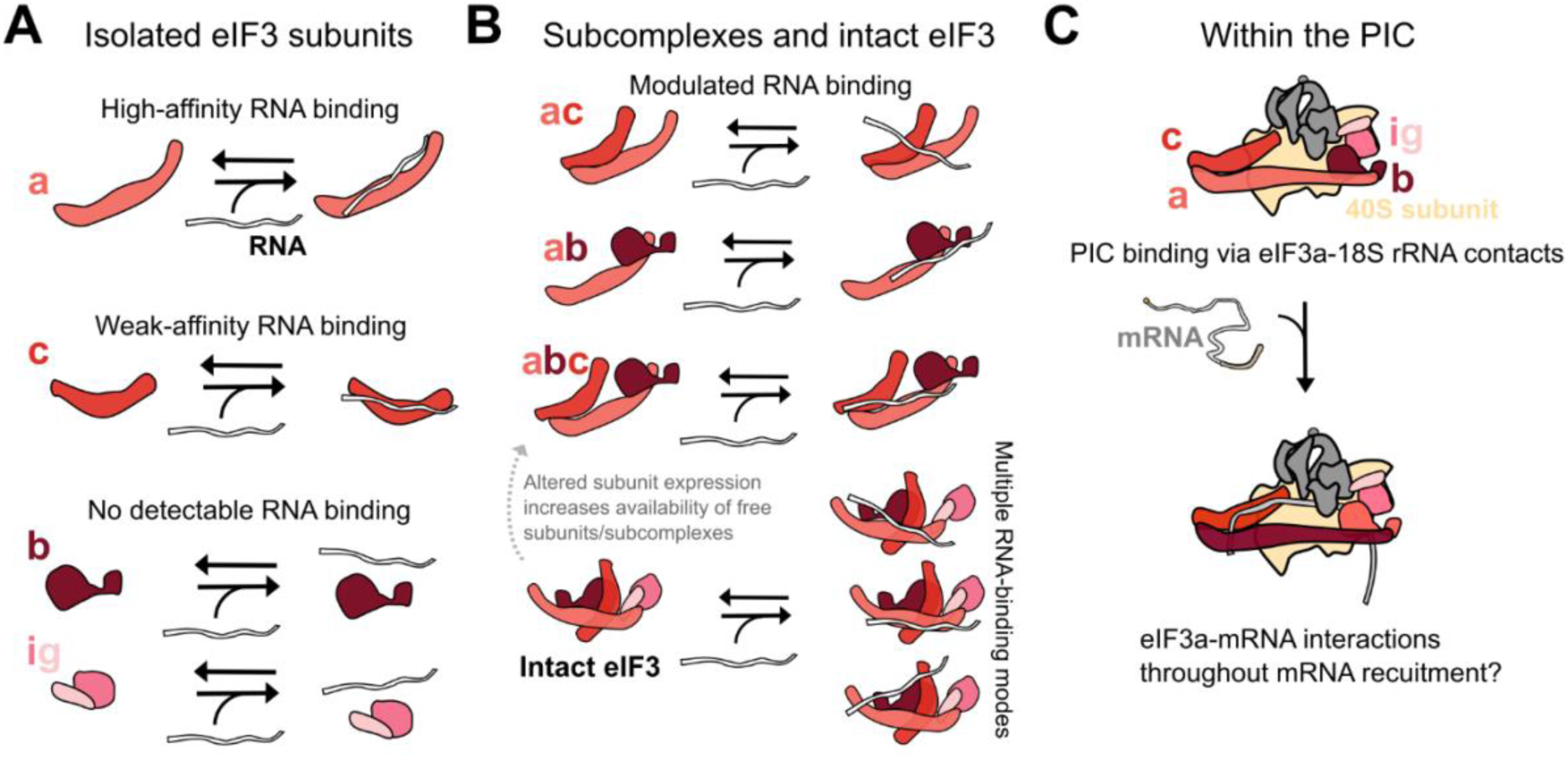
Subunit context tunes elF3a-driven RNA binding within elF3. Model summarizing the RNA-binding activities of individual elF3 subunits, defined elF3a-containing subcomplexes, and intact elF3. **(A)** Among the isolated subunits tested, elF3a provides the principal RNA-binding activity. elF3c binds RNA weakly, whereas elF3b and elF3ig show little or no detectable RNA binding under the conditions tested. **(B)** Defined elF3a-containing subcomplexes are competent to bind RNA, and the interaction between elF3a and elF3c alters the RNA-binding behavior of elF3a. Within intact elF3, interactions with other subunits may alter RNA contacts, the accessibility or geometry of the elF3a RNA-binding surface, or transitions among distinct RNA-bound states, producing multiple RNA-binding modes. Altered elF3 subunit abundance could also change the availability of free subunits or subcomplexes, although this possibility remains to be tested directly in cells. **(C)** Proposed deployment of elF3a-driven RNA binding during canonical initiation. elF3a may contribute initially to elF3 binding to the PIC through contacts with 18S rRNA. Following mRNA attachment to the PIC, the RNA-binding activity ofelF3a may be repurposed or remodeled to engage mRNA during loading, scanning, and start-codon recognition.

The tethered-RNA mass-photometry experiments provide complementary evidence that robust, repeated RNA-binding events depend strongly on eIF3a and that defined eIF3a-containing subcomplexes are competent to engage RNA. We did not use these experiments to infer the solution-phase composition, stability, or exchange behavior of eIF3. Instead, we used RNase sensitivity and filtering for repeated events as complementary operational criteria for identifying RNA-dependent apparent-mass distributions. These analyses showed that robust repeated events depend strongly on the presence of eIF3a. Experiments with separately prepared eIF3a, eIF3ac, eIF3ab, eIF3abc, and eIF3abig further showed that isolated eIF3a and multiple defined eIF3a-containing subcomplexes can engage tethered RNA. Because apparent-mass distributions cannot uniquely identify the molecular species producing events within an eIF3_R_ sample, these experiments do not establish the relative contributions of intact eIF3 and low-abundance subcomplexes to RNA binding by an eIF3_R_ preparation. Likewise, because event counts alone do not provide precise binding affinities, these experiments do not establish a thermodynamic rank ordering among intact eIF3 and the various eIF3a-containing subcomplexes. Nonetheless, they support the broader conclusion that eIF3a provides a core RNA-binding activity that is retained in multiple subunit contexts.

These findings raise the possibility that changes in eIF3 subunit abundance or assembly state could alter how eIF3 RNA-binding activity is deployed in cells. Under normal cellular conditions, the stoichiometry of eIF3 subunits in *S. cerevisiae* is expected to be nearly equal (von der Haar and McCarthy 2002; Ho, Baryshnikova and Brown 2018), such that intact five-subunit eIF3 is expected to be the predominant species. Changes in the abundance of individual eIF3 subunits could, in principle, alter the amounts of intact eIF3, free subunits, or subcomplexes with distinct biochemical activities. Consistent with the importance of maintaining appropriate subunit abundance, altered expression of individual yeast eIF3 subunits perturbed growth, with TEF1 - driven expression of eIF3a producing the strongest phenotype in our assays (Figure S7). These experiments do not establish whether these growth phenotypes result from changes in intact eIF3 assembly or abundance, accumulation of free subunits or subcomplexes, altered RNA binding, or other eIF3-dependent processes. They instead identify these possibilities as questions for future investigation. Because the five-subunit yeast eIF3 complex corresponds to the conserved core of the larger mammalian complex, this framework may also be relevant to human cells, where dysregulated expression of individual eIF3 subunits has been linked to disease-associated translational programs, including those associated with cancer (Zhang, Pan and Hershey 2007; Hershey 2015; Gomes-Duarte et al. 2018; Wolf et al. 2020).

The central role of eIF3a in RNA binding also has important implications for how eIF3 binds the PIC and supports mRNA attachment, loading, scanning, and start-codon recognition during mRNA recruitment. In the canonical initiation pathway, eIF3 binds the 40S subunit as part of the PIC, where eIF3a and other eIF3 subunits are positioned to contact the 18S rRNA (Figure 6C). Thus, eIF3a-driven RNA binding may contribute both to rRNA contacts that stabilize eIF3 binding to the PIC and to subsequent interactions with mRNA. Once eIF3 is bound to the PIC, the same RNA-binding activity could be repurposed or remodeled to engage the incoming mRNA during recruitment. One possible alternative model is that eIF3a-containing subcomplexes bind mRNA before engagement with the PIC and contribute directly to mRNA delivery. Although our data do not exclude this possibility, we favor a model in which the primary role of eIF3a-driven RNA binding during canonical initiation occurs within PIC-bound eIF3. In this model, initial delivery of activated mRNA to the PIC is mediated largely by the eIF4F-dependent mRNA-activation pathway (Gentry et al. 2025; Gentry et al. 2026), whereas eIF3a within PIC-bound eIF3 provides a tunable RNA-binding surface that contributes to PIC binding through rRNA contacts and facilitates subsequent steps of mRNA recruitment through contacts with mRNA (Figure 6C). This model is consistent with the high affinity of eIF3 for the PIC, the expected cellular concentrations of ribosomes and eIF3 in yeast, and prior work suggesting that PIC binding can allosterically modulate eIF3 structure (Zeman et al. 2019). eIF3a is particularly well positioned to coordinate these RNA interactions because PIC-bound eIF3a spans the mRNA-entry and mRNA-exit channel regions and provides a principal physical connection between eIF3 modules located at these sites (Erzberger et al. 2014; Llacer et al. 2015; Llacer et al. 2021).

More broadly, our results support a view of eIF3 as a modular RNA-binding platform whose activity is shaped by subunit context and interactions with the PIC. Within the conserved yeast core, eIF3a provides the principal RNA-binding activity, whereas other subunits modulate this activity and may create additional opportunities for allosteric control. In higher eukaryotes, additional eIF3 subunits, many of which are positioned near the mRNA-exit channel, may further expand the regulatory potential of this platform. These subunits may tune RNA engagement, promote mRNA-specific interactions, or couple eIF3 to regulatory pathways that alter initiation in response to cellular state. Defining how individual eIF3 subunits and subcomplexes contribute to RNA binding therefore provides a foundation for understanding how eIF3 binds the PIC, supports mRNA recruitment during canonical initiation, participates in alternative initiation pathways, and responds to regulated or dysregulated changes in subunit abundance and interactions.

## METHODS

### Cloning, expression, and purification of recombinant eIF3 subunits and subcomplexes

The expression plasmids for eIF3a and eIF3b were described previously (Zeman et al. 2019). The eIF3c expression plasmid encodes an N-terminal six-histidine (6× His) tag, a *Fasciola hepatica* 8-kDa (Fh8) solubility-enhancement tag, and a tobacco etch virus (TEV) protease recognition site upstream of full-length eIF3c. TEV cleavage yields untagged eIF3c without additional N-terminal residues. The plasmid encoding the eIF3i-eIF3g fusion was described previously (Zeman et al. 2019), except that the PreScission protease cleavage site between eIF3i and eIF3g was replaced with a TEV protease cleavage site.

Each recombinant subunit or fusion protein was expressed from a pET vector (Novagen) in *E. coli* BL21-CodonPlus (DE3)-RIPL cells (Agilent, Catalog Number 230280). Cultures were grown at 37 °C to an optical density at 600 nm (OD_600_) of 0.4–0.6, and the temperature was lowered to 17.5 °C. After 45–60 min, expression was induced with 0.5 mM isopropyl β-D-1-thiogalactopyranoside (IPTG), and cultures were grown for an additional 16–20 h. Cultures were cooled on ice, and cells were collected by centrifugation at 8,000 × g for 5 min. Cell pellets were resuspended in 40–50 mL of eIF3 Lysis Buffer (20 mM 4-(2-hydroxyethyl)-1-piperazineethanesulfonic acid (HEPES), pH 7.4, 400 mM potassium chloride (KCl), 10% glycerol, and 2 mM β-mercaptoethanol (BME)). The cell suspension was flash-frozen as droplets in liquid nitrogen. Frozen droplets were stored at −80 °C for up to 2 months before lysis in a freezer mill (SPEX, Catalog Number 6875D115) for seven cycles of 2 min at 10 cycles per second, with 1 min between cycles. The resulting lysate powder was stored at −80 °C.

For purification, lysate powder was slowly added to 100–200 mL of stirred, room-temperature eIF3 Lysis Buffer supplemented with two EDTA-free protease inhibitor tablets (Roche, Catalog Number 04693132001), 20 mM imidazole, 10 mM magnesium chloride (MgCl2), and 50 µg of nuclease A from *Serratia marcescens*. The lysate was clarified by centrifugation at 31,500 × g for 30 min, and the supernatant was passed through a 0.45-µm filter. The filtrate was loaded onto a 5-mL HiTrap Chelating column (Cytiva, Catalog Number 17040901) charged with nickel sulfate (NiSO_4_) according to the manufacturer’s instructions and equilibrated in K-200 Buffer (20 mM HEPES, pH 7.4, 200 mM KCl, and 10% glycerol). Bound protein was eluted with an eight-column-volume gradient of K-200 Buffer supplemented with 400 mM imidazole. The subsequent purification steps differed among proteins, as described below.

For eIF3a and eIF3b, the complete nickel-affinity eluate was pooled, diluted two-fold with K-200 Buffer, and loaded onto two 5-mL HiTrap Q columns connected in series (Cytiva, Catalog Number 17115401) and equilibrated in K-200 Buffer. Protein was eluted with a 15-column-volume gradient of K-1000 Buffer (20 mM HEPES, pH 7.4, 1,000 mM KCl, and 10% glycerol). Fractions were assessed by sodium dodecyl sulfate polyacrylamide gel electrophoresis (SDS-PAGE) and Coomassie brilliant blue staining. Fractions containing the desired protein were pooled, and protein concentration was determined by Bradford assay using bovine serum albumin as a standard. The proteins were divided into 10-µL single-use aliquots and flash-frozen in liquid nitrogen.

For eIF3c, nickel-affinity fractions containing eIF3c were pooled and treated with TEV protease at a protease-to-substrate ratio of 1:100 (w/w) in the presence of 2 mM BME. The mixture was incubated at room temperature for 2–3 h and then at 4 °C overnight. To reduce the imidazole concentration, the mixture was diluted ten-fold with K-200 Buffer and passed through a freshly NiSO_4_-charged 5-mL HiTrap Chelating column equilibrated in K-200 Buffer. The flow-through, containing cleaved eIF3c, was loaded onto two 5-mL HiTrap Q columns connected in series and eluted as described for eIF3a and eIF3b. Fractions were assessed by SDS-PAGE, and those containing eIF3c were pooled, divided into single-use aliquots, and flash-frozen in liquid nitrogen.

For the eIF3i-eIF3g fusion protein, nickel-affinity fractions containing the fusion protein were pooled, diluted twofold with K-200 Buffer, loaded onto two 5-mL HiTrap Q columns connected in series, and eluted as described above. Fractions containing the fusion protein were pooled and treated with TEV protease at a protease-to-substrate ratio of 1:100 (w/w) in the presence of 2 mM BME. The mixture was incubated at room temperature for 2–3 h and then at 4 °C overnight. Cleavage products were separated from TEV protease by size-exclusion chromatography on a Superdex 200 Increase 10/300 GL column (Cytiva, Catalog Number 28990944) equilibrated and run in K-200 Buffer. Fractions were assessed by SDS-PAGE; those containing detectable uncleaved fusion protein were excluded. Fractions containing co-purified eIF3i and eIF3g, referred to here as the eIF3ig subcomplex, were pooled. Protein concentration was determined, and the preparation was divided into single-use aliquots and flash-frozen in liquid nitrogen.

### Assembly of eIF3_R_ and defined subcomplexes

Unless otherwise indicated, eIF3_R_ was assembled by mixing purified eIF3a, eIF3b, eIF3c, and eIF3ig at 1:1:1:1 molar ratios and 2-5 µM total complex concentration in K-200 buffer. The mixture was incubated at room temperature for 20 minutes before use. Concentrations reported for eIF3_R_ refer to the concentration of each individual subunit.

Unless otherwise indicated, defined eIF3a-containing subcomplexes, including eIF3ac, eIF3ab, eIF3abc, and eIF3abig, were prepared by combining their purified components at equimolar ratios in K-200 buffer and incubating at room temperature for 20 minutes. For the eIF3ac anisotropy titration, the reported 5-µM concentration refers to the final concentrations of both eIF3a and eIF3c.

### Purification of eIF3_N_

eIF3N was purified from strain LPY87 as previously described (Acker et al. 2007; Liu et al. 2019), with the modifications specified below. LPY87 was grown on selective medium lacking uracil and leucine for 2–3 nights at 30 °C. A 50–100-mL starter culture was grown overnight in liquid selective medium at 30 °C. After 16–20 h, 5 mL of starter culture was added to 1.5 L of yeast extract-peptone-dextrose (YPD) medium and grown overnight at 30 °C. Cells were collected, resuspended, flash-frozen, and lysed in eIF3 Lysis Buffer as described above.

The lysate was loaded onto a 5-mL HiTrap Chelating column charged with NiSO_4_ and eluted as described for the recombinant subunits.

The complete nickel-affinity eluate was pooled, diluted twofold with K-200 Buffer, and loaded onto two 5-mL HiTrap Q columns connected in series. To seek separation of eIF3 complexes with differing amounts of co-purifying eIF5, a shallow 20-column-volume gradient from 0% to 50% K-1000 Buffer in K-200 Buffer was used (Llacer et al. 2021). Two major peaks eluted, one centered at a conductivity of 25 mS/cm and a broader peak at > 27 mS/cm. Fractions from each peak were assessed by SDS-PAGE and pooled separately. Each pool was then subjected to size-exclusion chromatography on a Superose 6 Increase 10/300 column (Cytiva, Catalog Number 29091596) equilibrated and run in K-200 Buffer. Each pool yielded a higher-molecular-mass and a lower-molecular-mass peak; the > 27-mS/cm pool contained a larger proportion of the higher-molecular-mass material. Fractions from the higher-molecular-mass peak of this pool were combined and assessed by SDS-PAGE for the five core eIF3 subunits. Protein concentration was determined by Bradford assay, and eIF3N was divided into 10-µL single-use aliquots and flash-frozen in liquid nitrogen.

### Purification of eIF1, eIF1A, eIF2, eIF4G, eIF4A, and eIF4B

eIF1, eIF1A, and eIF4A were expressed with N-terminal 6× His tags followed by TEV protease cleavage sites. eIF4B was expressed with a C-terminal TEV protease cleavage site followed by a 6× His tag. These factors were purified from *E. coli* BL21(DE3) RIPL cells (Agilent, Catalog Number 230280) using established procedures (Acker et al. 2007; Mitchell et al. 2010; Liu et al. 2019). Native 6× His-tagged eIF2 was expressed and purified from *S. cerevisiae* strain GP3511 as previously described (Acker et al. 2007). eIF4G bearing a C-terminal TEV protease cleavage site followed by an Fh8 tag and a 6× His tag was co-expressed with full-length eIF4E, yielding the eIF4G complex. This complex was purified from *E. coli* BL21(DE3) RIPL cells as previously described (Gentry et al. 2025).

### Purification of 40S subunits

Native *S. cerevisiae* 40S subunits were purified from the wild-type W303 strain LSY679 largely as described previously (Acker et al. 2007), with the modifications specified below. Cells were grown at 30 °C to an OD_600_ of 0.8–1.2, collected, resuspended in Ribosome Lysis Buffer (20 mM HEPES, pH 7.4, 100 mM potassium acetate (KOAc), 7.5 mM magnesium acetate (Mg(OAc)_2_), and 5 mM BME), and flash-frozen as droplets in liquid nitrogen. Frozen droplets were lysed in a freezer mill as described above.

Lysate powder was resuspended in 50–200 mL of room-temperature Ribosome Lysis Buffer supplemented with two EDTA-free protease inhibitor tablets (Roche, Catalog Number 04693132001), TURBO DNase (Invitrogen, Catalog Number AM2239), and 0.5–1 mg/mL heparin. The lysate was clarified at 31,500 × g for 30 min. High-Salt Sucrose Cushion Buffer (20 mM HEPES, pH 7.4, 100 mM KOAc, 500 mM KCl, 2.5 mM Mg(OAc)_2_, 34% (w/v) sucrose, and 5 mM BME) was layered beneath the clarified lysate. Ribosomes were pelleted through the cushion at 60,000 rpm in a Ti 70 rotor (Beckman Coulter, Catalog Number 337922) for 2 h at 4 °C. The pellets were rinsed with 1 mL of Subunit Separation Buffer (50 mM HEPES, pH 7.4, 550 mM KCl, 2 mM MgCl_2_, and 5 mM BME). Tubes were inverted for 10 min at room temperature, and the pellets were resuspended in approximately 400 µL of Subunit Separation Buffer.

Resuspended ribosomes were pooled, and puromycin dihydrochloride (GoldBio, Catalog Number P-600-100) was added to 2 mM to promote dissociation of 40S and 60S subunits. The mixture was incubated on ice for 10 min and at 37 °C for 15 min, then layered onto 10%–35% sucrose density gradients prepared in Subunit Separation Buffer. Subunits were separated by ultracentrifugation in an SW32 rotor (Beckman Coulter, Catalog Number 369694) at 22,000 rpm for 17 h at 4 °C.

Gradients were fractionated using a BioComp Instruments Piston Gradient Fractionator. Fractions corresponding to 40S and 60S subunits were pooled separately and pelleted through High-Salt Sucrose Cushion Buffer as described above. Each pellet was resuspended in Subunit Separation Buffer and subjected to a second 10%–35% sucrose density gradient to reduce cross-contamination of the subunits. The 40S and 60S subunit fractions were again collected separately and pelleted through Ribosome Storage Cushion Buffer (20 mM HEPES, pH 7.4, 100 mM KOAc, 2.5 mM Mg(OAc)_2_, 30% (w/v) sucrose, and 2 mM dithiothreitol (DTT)). Pellets were resuspended in Ribosome Storage Buffer (20 mM HEPES, pH 7.4, 100 mM KOAc, 2.5 mM Mg(OAc)_2_, 8.5% (w/v) sucrose, and 2 mM DTT). Concentrations were determined from absorbance at 260 nm using extinction coefficients of 2 × 10⁷ M⁻¹ cm⁻¹ and 4 × 10⁷ M⁻¹ cm⁻¹ for 40S and 60S subunits, respectively. Samples were divided into 5-µL single-use aliquots, flash-frozen in liquid nitrogen, and stored at −80 °C.

### Transcription, purification, and aminoacylation of initiator tRNA

*S. cerevisiae* initiator tRNA (tRNA_i_^Met^) was generated by *in vitro* transcription with T7 RNA polymerase. A DNA template containing a T7 promoter, a hammerhead ribozyme, and the tRNA_i_ sequence was amplified from a pUC19-derived plasmid by polymerase chain reaction. The hammerhead ribozyme enabled efficient transcription from a 5′ guanosine and subsequent production of tRNA with a native 5′ adenosine following self-cleavage of the transcript at 65 °C for 1 h. Cleavage products were separated by denaturing PAGE on a 7% polyacrylamide (19:1 acrylamide) gel prepared in TBE Buffer (90 mM Tris-borate and 2 mM EDTA) with 7 M urea. The tRNA band was identified by ultraviolet shadowing, excised, and extracted by overnight agitation at 4 °C in 300 mM sodium acetate, pH 5.2, with 1 mM EDTA. The extracted tRNA was concentrated by ethanol precipitation.

The tRNA pellet was dissolved in 10 mM sodium acetate, pH 5.2. To fold the tRNA, the solution was placed in boiling water and allowed to cool slowly to room temperature; 5 mM Mg(OAc)_2_ was added when the solution approached room temperature. Folded tRNA was aminoacylated in a reaction containing 100 mM HEPES, pH 7.4, 10 mM MgCl_2_, 10 mM KCl, 0.1 mM methionine, 10 mM ATP·Mg, 1 mM DTT, 1 µM *E. coli* methionyl-tRNA synthetase, and 12 µM folded tRNA ^Met^, based on a previously described procedure (Walker and Fredrick 2008). After 30 min at 37 °C, the mixture was phenol-chloroform extracted and ethanol precipitated. To remove residual ATP, the pellet was dissolved in 10 mM sodium acetate, pH 5.2, and passed sequentially through three PD SpinTrap G-25 columns (Cytiva, Catalog Number 28918004) equilibrated in the same buffer. Met-tRNA ^Met^ concentration was determined from absorbance at 260 nm using an extinction coefficient of 708,900 M⁻¹ cm⁻¹. The sample was divided into 5-µL single-use aliquots, flash-frozen in liquid nitrogen, and stored at −80 °C.

### Fluorophore labeling of eIF3b

The single-cysteine eIF3b^S460C^ variant was purified as described for wild-type eIF3b. Janelia Fluor 549 (JF549; Tocris, Catalog Number 6500) was stored as a lyophilized powder at −20 °C. For labeling, JF549 was dissolved to 10 mM in fresh anhydrous dimethyl sulfoxide, added in five-fold molar excess to eIF3b^S460C^, and incubated for 2 h at room temperature. The reaction was quenched with 2 mM DTT. Unreacted fluorophore was removed by size-exclusion chromatography on a Superdex 200 Increase 10/300 GL column (Cytiva, Catalog Number 28990944) equilibrated and run in K-200 Buffer. Fractions containing labeled (JF549)eIF3b were pooled, divided into 10-µL single-use aliquots, and flash-frozen in liquid nitrogen.

### Standard Reaction Buffer

Unless otherwise indicated, biochemical measurements were performed in Standard Reaction Buffer (20 mM HEPES, pH 7.4, 100 mM KOAc, and 3 mM Mg(OAc)_2_), which supports *in vitro S. cerevisiae* translation-initiation assays.

### Stability and exchange experiments using fluorophore-labeled eIF3b

For dilution experiments, a (JF549)eIF3b-containing eIF3R complex was assembled by combining 2 µM each of eIF3a, (JF549)eIF3b, eIF3c, and eIF3ig and incubating for 20 min at room temperature. The complex was diluted in Standard Reaction Buffer to concentrations of 30– 300 nM, as indicated. Within approximately 3 min of dilution, 800 µL of sample was loaded onto a Superose 6 Increase 10/300 column (Cytiva, Catalog Number 29091596). The eluate was collected as 200-µL fractions in 96-well plates (Corning, Catalog Number 3916). Fluorescence was measured on a BioTek Synergy Neo2 plate reader (Agilent) using a filter cube with 530-nm excitation and 590-nm emission filters (Agilent, Catalog Number 1505004).

For exchange experiments, unlabeled eIF3R was assembled by combining 2 µM each of eIF3a, eIF3b, eIF3c, and eIF3ig and incubating for 20 min at room temperature. The complex was diluted to 1 µM and mixed with 0.2 µM (JF549)eIF3b or (JF549)eIF3big to initiate exchange. After incubation at room temperature for the indicated times, 100 µL of each reaction was loaded onto a Superose 6 Increase 10/300 column. Fractions were collected, and fluorescence was measured as described above.

### PIC binding assays

Binding of eIF3_N_ and eIF3_R_ to PICs was assessed by native gel electrophoresis using a 3′-fluorescein-labeled polyCAA model mRNA with the sequence shown under ‘RNA binding assays’. A 2× PIC master mix contained 2 µM eIF1, 2 µM eIF1A, 600 nM eIF2, 750 nM Met-tRNA_i_ ^Met^, 800 nM GDPNP•Mg2+, 100 nM 40S subunits, and 1 µM labeled polyCAA RNA. The master mix was combined with eIF3_N_ or eIF3_R_ and Standard Reaction Buffer to yield final concentrations of 1 µM eIF1, 1 µM eIF1A, 300 nM eIF2, 375 nM Met-tRNA_i_ ^Met^, 400 nM GDPNP•Mg2+, 50 nM 40S subunits, 0.5 µM labeled polyCAA RNA, and 10–800 nM eIF3N or eIF3_R_. Reactions were incubated at 26 °C for 1 h. An 8-µL aliquot was mixed with 1 µL of Native Loading Dye (0.05% bromophenol blue and 50% sucrose in Standard Reaction Buffer) and loaded onto a 4% polyacrylamide (37.5:1 acrylamide) gel prepared in THEM Buffer (34 mM Tris base, 57 mM HEPES, 0.1 mM EDTA, and 2.5 mM MgCl_2_). The gel was run at 200 V for 1 h while maintained at 16 °C and imaged for fluorescein fluorescence on an Amersham Typhoon 5 Biomolecular Imager (Cytiva, Catalog Number 29187191). Because labeled RNA was present in excess over 40S subunits, the free-RNA signal saturated the detector and was excluded from PIC-binding quantification, as described below.

### mRNA recruitment assays

The kinetics of capped rpl41a mRNA recruitment to PICs were measured by a previously described native gel assay using [^32^P]-cap-labeled rpl41a mRNA (Mitchell et al. 2010; Aitken et al. 2016; Yourik et al. 2017). A 2× PIC master mix contained 2 µM eIF1, 2 µM eIF1A, 600 nM eIF2, 750 nM Met-tRNA ^Met^, 800 nM GDPNP•Mg2+, 100 nM eIF4G, 4 µM eIF4A, 800 nM eIF4B, 100 nM 40S subunits, and 4 mM ATP·Mg²⁺. eIF3_N_ or eIF3_R_ was added at the indicated final concentrations ranging between 50 and 400 nM, and reactions were initiated with 15 nM [^32^P]-cap-labeled rpl41a mRNA. Reactions were incubated at 26 °C. At the indicated times, 4 µL of reaction was mixed with 1 µL of Native Loading Dye and loaded onto an actively running 4% polyacrylamide (37.5:1 acrylamide) gel prepared in THEM Buffer. The gel was run at 200 V for 70 min while maintained at 16 °C, exposed to a phosphor screen overnight, and imaged on an Amersham Typhoon 5 Biomolecular Imager. Formation of the gel-shifted complex in this assay depends on start-codon recognition and therefore reports completion of the mRNA-recruitment process, including mRNA attachment and loading, scanning, and start-codon recognition.

### RNA binding assays

Direct RNA binding was assessed by fluorescence anisotropy using chemically synthesized, 3′-fluorescein-labeled model RNAs (Integrated DNA Technologies). The substrate sequences, written 5′ to 3′, were:

- polyCAA (50 nt): GGCAACAACAACAACAACAACAAAUGGAACAACAACAACAACAAAGUCGA–fluorescein
- polyCAA (45 nt): GGCAACAACAACAACAACAAAUGGAACAACAACAACAACAGUCGA-fluorescein
- polyCAA (40 nt): GGCAACAACAACAACAACAAAUGGACAACAACAAAGUCGA-fluorescein
- polyUC (41 nt): GGAAUCUCUCUCUCUCUCUAUGCUCUCUCUCUCUCUCUCUC–fluorescein
- *rpl41a*-derived (51 nt): GGAGACCACAUCGAUUCAAUCGAAAUGAGAGCCAAGUGGAGAAAGAAGAGA–fluorescein

For affinity measurements, Standard Reaction Buffer containing 5 nM labeled RNA was prepared. The highest-concentration protein sample was also adjusted to 5 nM labeled RNA, then serially diluted twofold with the 5-nM RNA solution to maintain a constant labeled-RNA concentration. Maximum protein concentrations were 800 nM for eIF3_N_ or eIF3_R_, 2 µM for isolated subunits or eIF3ig, and 5 µM for eIF3ac. Extended eIF3c titrations reached 5 µM where indicated. Minimum concentrations were 3.1 nM for eIF3_N_ or eIF3_R_ and 7.8 nM for isolated subunits or eIF3ig; the minimum eIF3ac concentration was 4.9 nM. Samples were transferred to a black 96-well plate (Corning, Catalog Number 3916), protected from light, and equilibrated for 20 min at room temperature. Fluorescence polarization was measured on a BioTek Synergy Neo2 plate reader (Agilent) using 485-nm excitation and an Agilent Green FP filter cube (Catalog Number 8040561).

For titration-regime experiments, total polyCAA RNA concentrations were 400, 600, or 800 nM, with 50 nM labeled RNA and 350, 550, or 750 nM unlabeled RNA, respectively. Protein titrations and equilibration were performed as above. For Figure 4, measured anisotropy values were converted to fractional saturation using min-max normalization within each replicate. Mean fractional saturation and error bars were calculated as the mean and standard deviation of these values, respectively.

### Yeast spotting assays

The centromeric origin of replication in the *E. coli-S. cerevisiae* shuttle vector pRG216 (Addgene Catalog Number 64528) was replaced with a multicopy 2-micron origin. Individual eIF3 subunit coding sequences were inserted under either a high-expression TEF1 promoter/terminator pair or a lower-expression RNR2 promoter/CYC1 terminator pair. Plasmids were introduced into wild-type *S. cerevisiae* W303 strain LSY679 by chemical transformation and selection on medium lacking uracil. Transformants were grown with shaking for 2 days at 30 °C in selective medium. Cultures were normalized to an OD_600_ of 0.5, serially diluted five-fold for five or six steps, and 2 µL of each dilution was spotted onto selective medium lacking uracil. Plates were incubated at 30 °C for 3 days before imaging (Figure S7).

### Mass photometry assays

Conventional mass photometry was performed on a TwoMP instrument (Refeyn). Borosilicate coverslips (Research Products International, Catalog Number 246008) were cleaned by sonication in 100% ethanol for 15 min and 1 M KOH for 15 min, rinsed thoroughly with nanopure water, and dried with pre-purified argon. Silicone gaskets (Grace Bio-Labs, Catalog Number GBL103250) formed sample wells. Proteins were diluted in Standard Reaction Buffer to 30 nM unless otherwise indicated. eIF3_N_ and eIF3_R_ were also measured at 1–30 nM after either 20 min of equilibration at room temperature or immediate dilution from micromolar concentrations. Measurements were acquired as 60-s movies with the regular 10.9-µm × 4.3-µm field of view in AcquireMP and analyzed in DiscoverMP (Refeyn). Apparent masses were calibrated using conalbumin (75 kDa), aldolase (158 kDa), and thyroglobulin (669 kDa) from a High Molecular Weight Calibration Kit (Cytiva, Catalog Number 28-4038-42). Because these measurements detect molecules through interactions with the imaging surface and yielded apparent-mass distributions inconsistent with fluorescence-tracked size-exclusion and exchange measurements, they were not used to infer solution-phase eIF3 composition, stability, or subunit exchange.

For tethered-RNA mass photometry, borosilicate coverslips were passivated with a mixture of methoxy-terminated PEG (Laysan Bio, Catalog Number MPEGSIL50001G) and biotin-terminated PEG (Laysan Bio, Catalog Number Biotin-PEG-SIL-5K-1g) as described previously (Gentry et al. 2025). Sample wells were formed by epoxying cups cut from P200 pipette tips to the derivatized surface (Devcon Home, Catalog Number 20945), which prevented leakage beneath silicone gaskets on the hydrophilic surface. Wells were rinsed with 70 µL of Standard Reaction Buffer, 70 µL of Standard Reaction Buffer containing 0.1% Tween-20, and then three times with 70 µL of Standard Reaction Buffer. Streptavidin (20 µL of 10 nM; Invitrogen, Catalog Number S888) was added, and unbound streptavidin was removed with four 70-µL rinses of Standard Reaction Buffer. A 3′-biotinylated version of the polyCAA RNA was tethered by adding 10 nM RNA for 1 min at room temperature, followed by five 70-µL buffer rinses to remove unbound RNA.

For each tethered-RNA measurement, 20 µL of 2× protein solution was mixed with 20 µL of Standard Reaction Buffer in the well and immediately imaged. Final concentrations were 200 nM eIF3_R_ or 200 nM of each specified protein species unless otherwise indicated. For the eIF3bcig condition, eIF3b, eIF3c, and eIF3ig were added such that each of the four constituent subunits was present at 200 nM. Separate measurements used eIF3a, eIF3ac, eIF3ab, eIF3abc, or eIF3abig at 200 nM. Ten consecutive 15-s movies were acquired with the regular field of view. Nuclease A from *Serratia marcescens* (10 ng) was then added to the well and incubated for 2 min before acquisition of ten additional 15-s movies. Multiple short movies were used because heating by the detection laser could perturb the PEG-derivatized surface. These experiments were used to identify apparent-mass classes of repeated, RNA-dependent events, not to measure solution-phase composition, subunit exchange, or relative binding affinities.

### Mass photometry data processing

Events detected by DiscoverMP were exported as comma-separated value files. Adsorption events assigned positive apparent masses and desorption events assigned negative apparent masses were retained in the exported data. Histograms displayed in Figure S3 were calculated from adsorption events with apparent masses between 0 and 1000 kDa using 5-kDa bins. Where normalized distributions are shown, bin counts were divided by the total number of included events so that bin frequencies summed to one. Where event counts are shown, counts were not normalized. Source event files are available as comma-separated values (CSV) files.

For tethered-RNA experiments, to identify repeated events at a given position, super-resolution x and y coordinates were rounded to the nearest integer pixel, and the number of events at each pixel was counted across the movie before changing the field-of-view. This was repeated for each of the ten 15-s movies. Distributions were then filtered for pixels with more than zero, one, two, or three events, as indicated in Figure 5E or 5F. This operational filter enriches for repeated association and dissociation at tethered RNA but does not uniquely identify a bound protein species.

### Quantification of PIC binding and mRNA recruitment assays

TIFF images from the PIC-binding and mRNA-recruitment assays were analyzed using ImageQuant TL, version 8.2.0.0 (Cytiva). In PIC-binding gels, bands corresponding to free RNA, RNA·PIC, and RNA·PIC·eIF3 were identified. After minimum-profile background subtraction, the fraction of eIF3-bound PICs was calculated as the intensity of the RNA·PIC·eIF3 band divided by the sum of the RNA·PIC and RNA·PIC·eIF3 band intensities; the saturated free-RNA band was excluded. For mRNA-recruitment gels, rubber-band background correction was applied, and the fraction of recruited mRNA was calculated as the intensity of the mRNA·PIC band divided by the total lane intensity. Recruitment time courses were fit by nonlinear least squares using Python’s curve_fit function to a single-exponential association model, *f*(*t*) = *A*(1 − *e*^−*k*^*^obs^*^∗^*^t^*), where *f(t)* is the fraction of recruited mRNA at time *(t)*, *A* is the plateau fraction, and *k_obs_* is the observed rate constant.

### Determination of equilibrium dissociation constants

Binding data were fit to the quadratic binding equation:

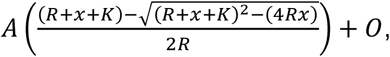

where *y* is the measured response, *A* is the binding-dependent amplitude, *R* is the total concentration of labeled RNA, *x* is the total concentration of the titrated protein species, *K* is *K*_d_^app^, and *O* is the response offset. PIC-binding data were fit the same way, with *R* representing total PIC concentration (50 nM) and *x* representing total eIF3 concentration. All fitting was done by nonlinear least squares using Python’s curve_fit function.

To model expected titration-regime behavior at increasing total RNA concentrations for eIF3R and eIF3a (Figure 4), *K* was fixed at the experimentally measured mean *K*_d_^app^ for each protein species, *O* was fixed at zero, and *A* was fixed at one. The predicted fractional-saturation curves were compared directly with the normalized experimental data without further scaling. These model curves represent predictions of a two-component, single-site interaction and were not used to assign a unique *K*_d_^app^ to the non-monotonic eIF3_R_ titrations

## ACKNOWLEDGEMENTS

This work was supported by funds to R.L.G. and C.E.A. from the NIH (R01 CA277727, R35 GM153724, and R01 GM084288 to R.L.G. and R15 GM140372 to C.E.A.). R.C.G. and N.A.I. were supported by NSF GRFP fellowships (DGE-1644869 and DGE-2036197, respectively). E.W.H. was funded through an F32 fellowship from the NIH (F32 GM139360). The authors would like to thank Dr. Eric Greene for his thoughtful and critical suggestions on early versions of the manuscript, the Columbia Precision Biomolecular Characterization Facility (PBCF) for access to instrumentation, and the PBCF Manager, Dr. Jerry Chang, for assistance with instrumentation.

## AUTHOR CONTRIBUTIONS

Conceptualization, N.A.I., R.C.G., C.D.K.-T., R.L.G., and C.E.A.; Methodology, N.A.I., R.C.G., C.D.K.-T., R.L.G., and C.E.A.; Investigation, N.A.I., R.C.G., B.A., and M.A.R.; Resources, N.A.I., R.C.G., V.M.C., K.Y., B.A., and E.W.H.; Validation, N.A.I., R.C.G., M.A.R., B.A., and P.K.V.; Writing – Original Draft, N.A.I., R.C.G., R.L.G., and C.E.A.; Writing – Review & Editing, N.A.I., R.C.G., R.L.G., and C.E.A.; Supervision, R.L.G. and C.E.A.; Funding acquisition, R.L.G. and C.E.A.

## MATERIALS AND CORRESPONDENCE

Further information and requests for resources and reagents should be directed to and will be fulfilled by either of the corresponding authors, Ruben L. Gonzalez, Jr. or Colin Echeverría Aitken.

## SUPPLEMENTARY FIGURES

**Supplemental Figure 1.**
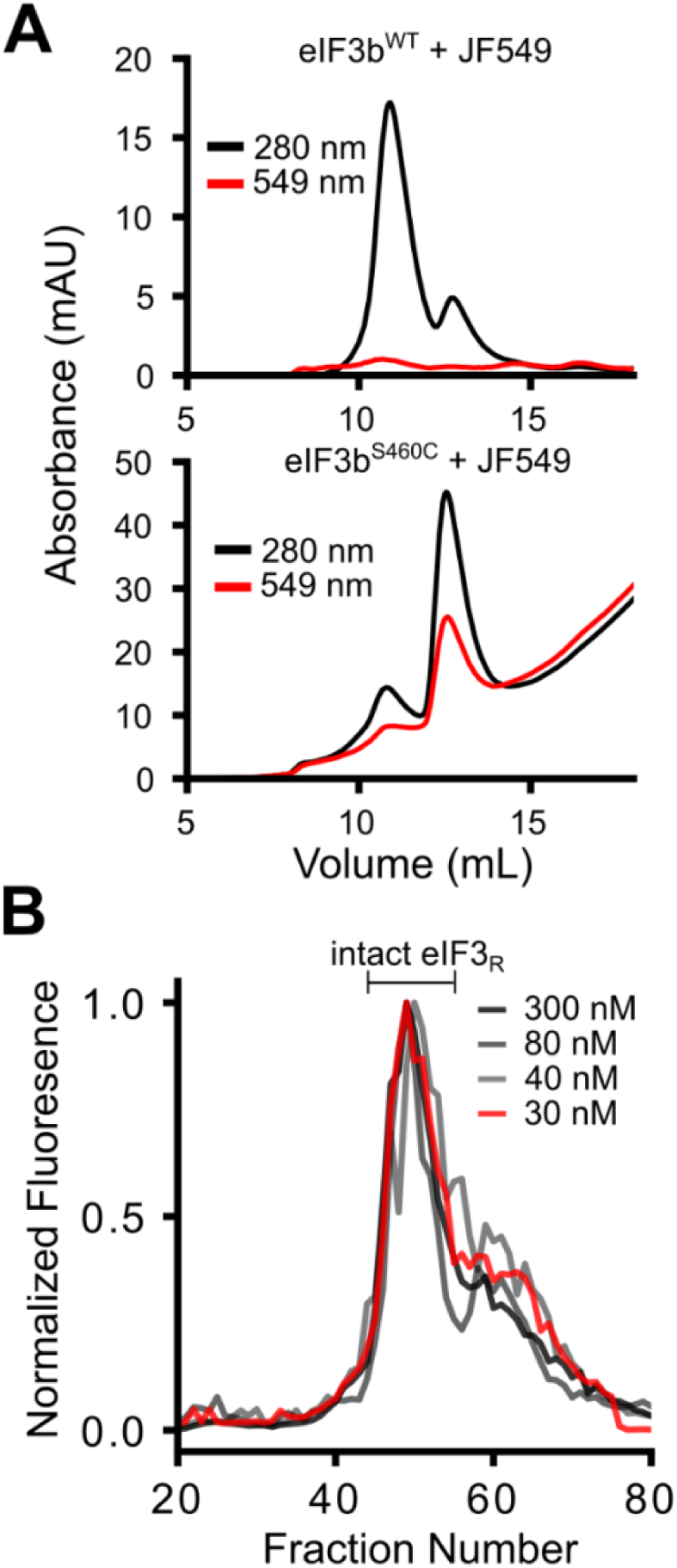
Selective fluorophore labeling of eIF3b^S460C^ with JF549 and stability of diluted (JF549)eIF3b-containing eIF3_R_. **(A)** Absorbance size-exclusion chromatograms of wild-type eIF3b and eIF3b^S460C^ following incubations with JF549. Absorbance at 280 nm reports on protein, and absorbance at 549 nm reports on JF549. The substantially greater co-elution of the 549-nm signal with eIF3bS460C than with wild-type eIF3b supports preferential labeling of the engineered cysteine. Excess free fluorophore is not shown. **(B)** Fluorescence size-exclusion chromatograms of (JF549)eIF3b-containing eIF3_R_ following dilution to the indicated concentrations. Fluorescence profiles were normalized to the maximum signal in each chromatogram. Across the tested concentration range, fluorescence remained detectable in fractions corresponding to intact eIF3_R_, consistent with retention of (JF549)eIF3b within the complex under these conditions.

**Supplemental Figure 2.**
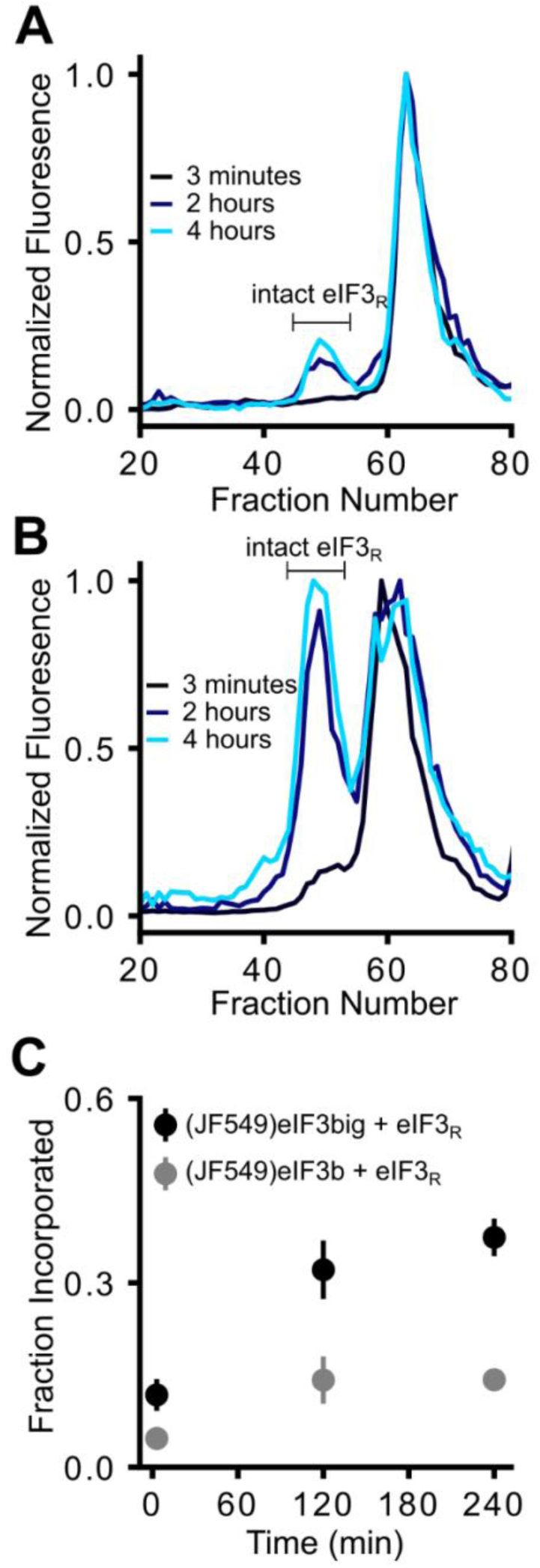
Slow exchange of (JF549)eIF3b and (JF549)eIF3big into intact eIF3_R_. **(A)** Fluorescence size-exclusion chromatograms of reactions containing 0.2 µM (JF549)eIF3b and 1 µM pre-formed, unlabeled eIF3_R_ after incubation for the indicated times. **(B)** Fluorescence size-exclusion chromatograms of reactions containing 0.2 µM (JF549)eIF3big and 1 µM pre-formed, unlabeled eIF3_R_ after incubation for the indicated times. Fluorescence profiles in panels A and B were normalized to the maximum signal in each chromatogram. Fractions corresponding to intact eIF3_R_ are indicated. **(C)** Fraction of total fluorescence recovered in fractions corresponding to intact eIF3_R_ for the experiments shown in panels A and B. Points represent the mean of two independent experiments, and error bars represent the standard deviation. Incorporation of both labeled species increased over hours, with (JF549)eIF3big incorporating more rapidly and to a greater extent than (JF549)eIF3b alone. These results indicate that exchange into intact eIF3_R_ is slow relative to the timescale of the biochemical assays

**Supplemental Figure 3.**
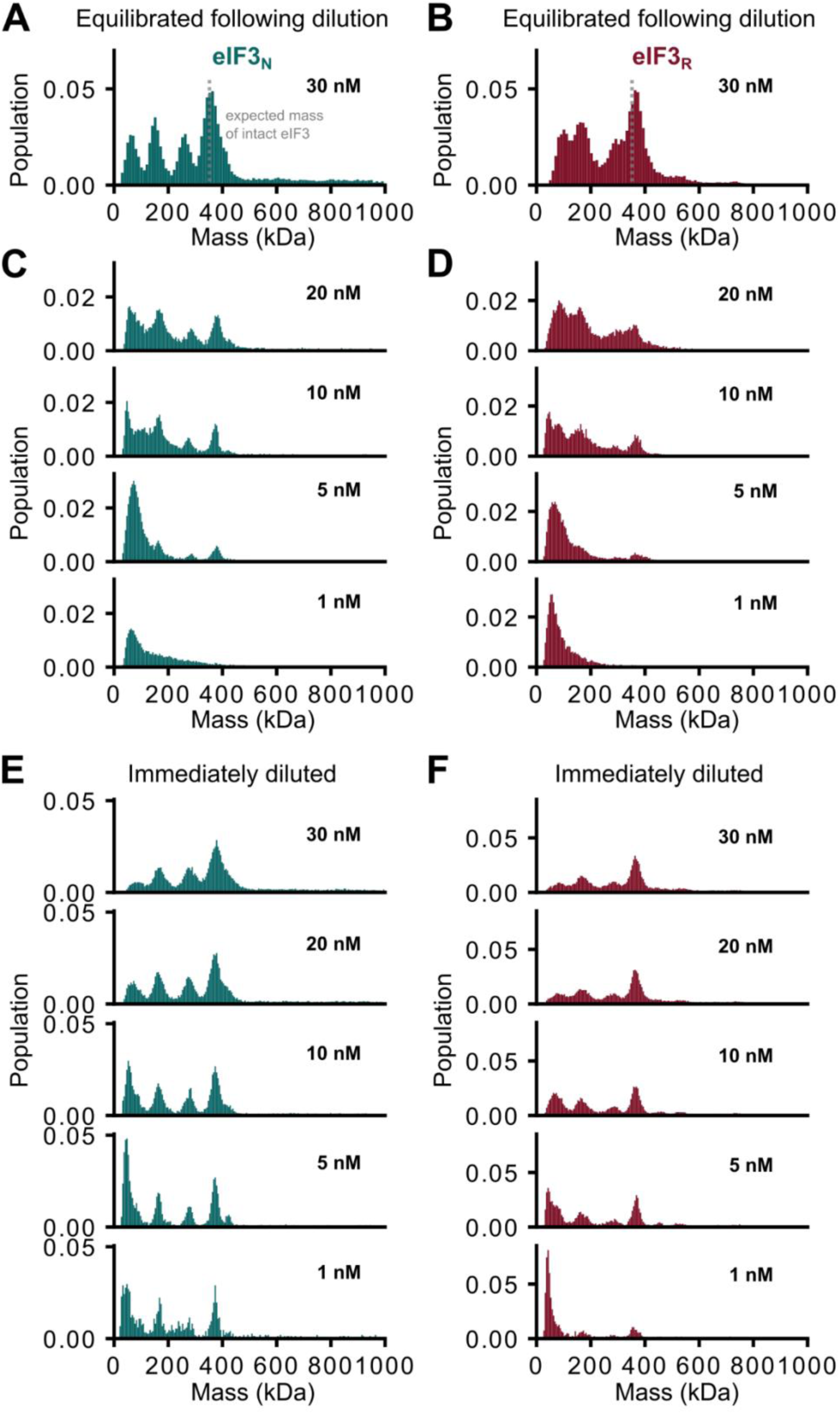
Conventional mass photometry of eIF3_N_ and eIF3_R_ yields multiple surface-associated apparent-mass classes at low nanomolar concentrations. **(A, B)** Apparent-mass distributions obtained by conventional mass photometry for 30 nM (A) eIF3_N_ or **(B)** eIF3_R_ after equilibration for at least 20 min following dilution from micromolar concentrations. **(C, D)** The expected molecular mass of intact eIF3, ∼360 kDa, is indicated by a dashed gray line for reference and is not intended as an assignment of an observed population. Apparent-mass distributions obtained for **(C)** eIF3_N_ or **(D)** eIF3_R_ at the indicated concentrations after equilibration for at least 20 min following dilution. **(E, F)** Apparent-mass distributions obtained for **(E)** eIF3_N_ or **(F)** eIF3_R_ immediately after dilution from micromolar concentration. Distributions were normalized as described in the Methods. These measurements report on detected surface-associated events and were not used to infer solution-phase eIF3 composition, stability, or subunit-exchange behavior.

**Supplemental Figure 4.**
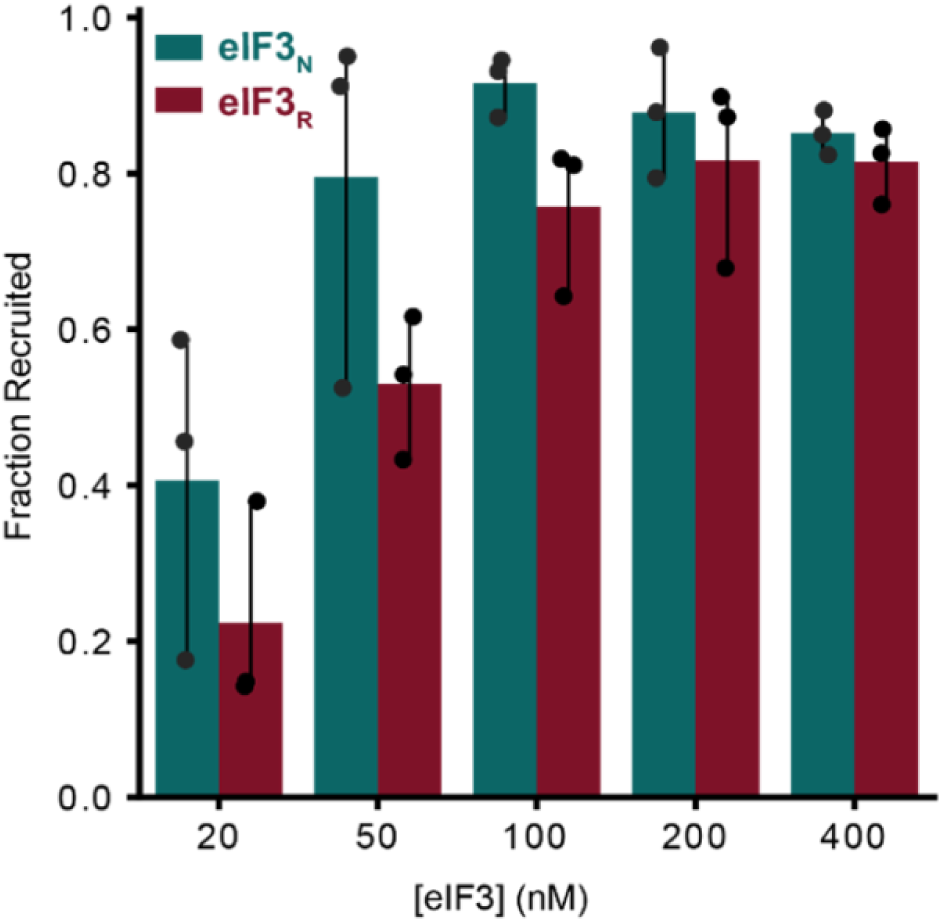
Concentration-dependent stimulation of capped rpl41a mRNA recruitment by eIF3_N_ and eIF3_R_. Fraction of capped rpl41a mRNA recruited into PICs after 60 min in the presence of the indicated concentrations of eIF3_N_ or eIF3_R_. Bars represent the mean of three independent experiments, individual replicate values are shown as points, and the standard deviation is represented by the error bars. Both eIF3_N_ and eIF3_R_ stimulated mRNA recruitment in a concentration-dependent manner and produced similarly high recruitment endpoints at concentrations of 100 nM or greater.

**Supplemental Figure 5.**
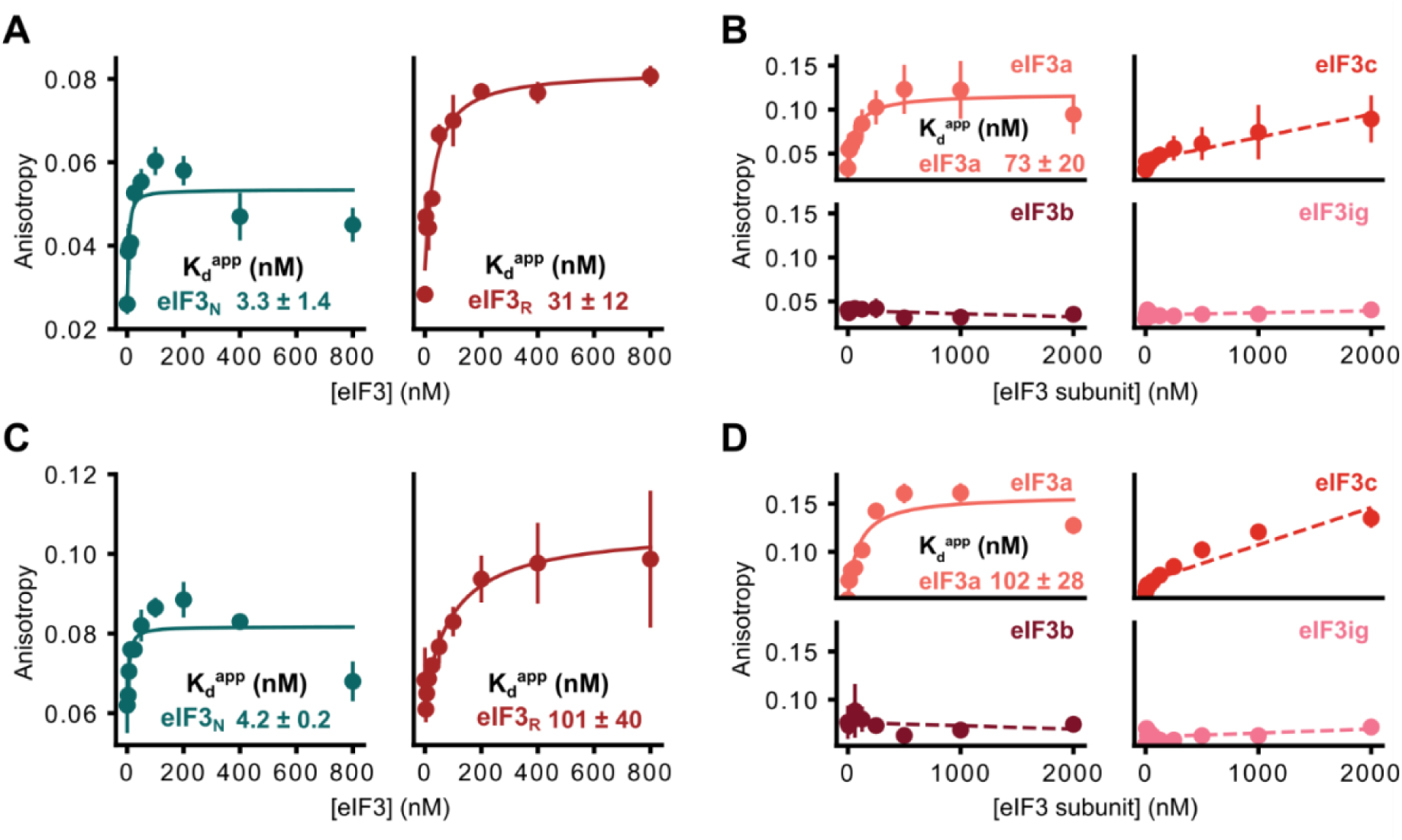
RNA binding by eIF3_N_, eIF3_R_, and isolated eIF3 subunits across sequence-distinct RNAs. **(A)** Fluorescence-anisotropy measurements of eIF3_N_ and eIF3_R_ binding to fluorescein-labeled polyUC RNA. **(B)** Fluorescence-anisotropy measurements of isolated eIF3a, eIF3b, eIF3c, and the co-purified eIF3ig subcomplex binding to polyUC RNA. **(C)** Fluorescence-anisotropy measurements of eIF3_N_ and eIF3_R_ binding to fluorescein-labeled rpl41a-derived RNA. **(D)** Fluorescence-anisotropy measurements of isolated eIF3a, eIF3b, eIF3c, and eIF3ig binding to rpl41a-derived RNA. Points represent the mean of three independent titrations, and error bars represent the standard deviation. Solid curves show fits to the quadradic binding equation. Dashed lines are included only as visual guides for titrations that did not approach saturation and do not represent reliable binding-model fits. Together with the polyCAA measurements shown in Figure 3, these data show that eIF3_N_, eIF3_R_, and isolated eIF3a bind each of the three tested RNA sequences without evidence of a consistent sequence preference.

**Supplemental Figure 6.**
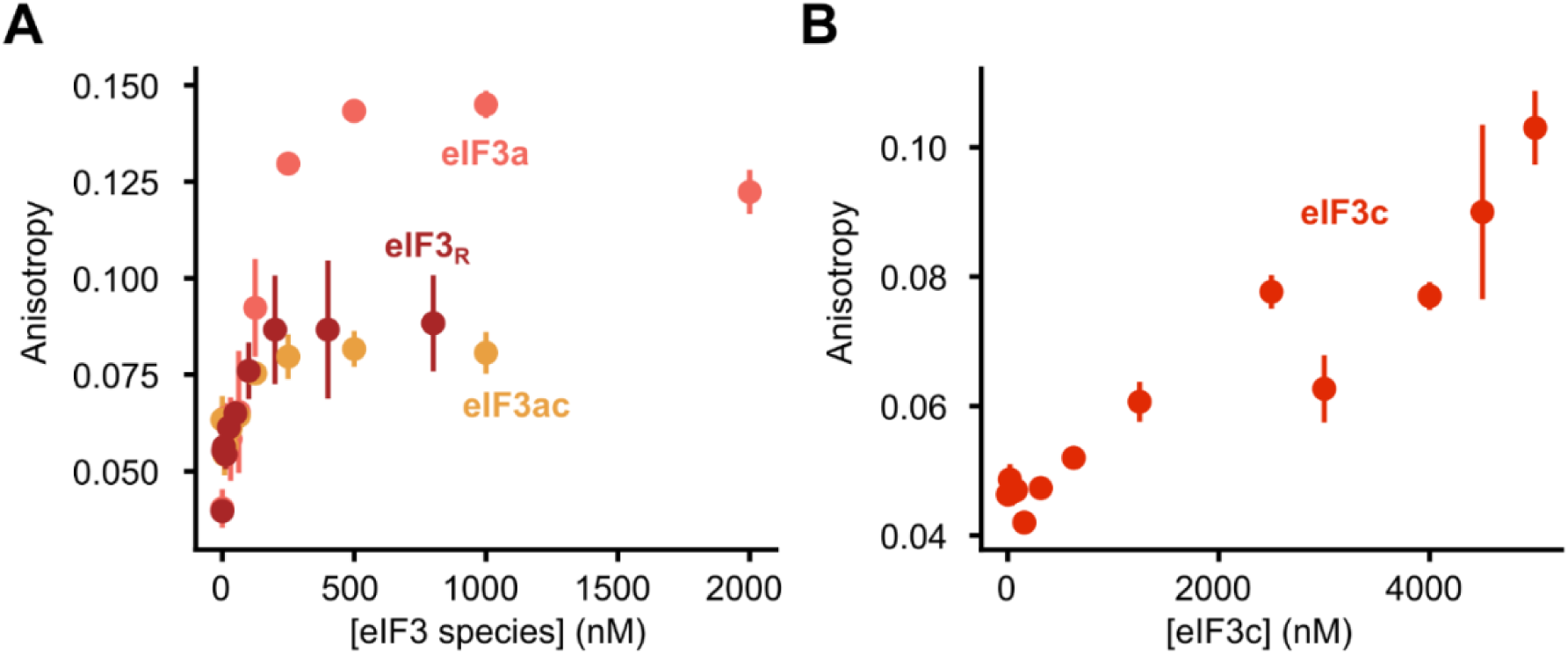
RNA binding by eIF3ac and extended eIF3c titration. **(A)** Fluorescence-anisotropy measurements of eIF3a, eIF3_R_, and eIF3ac binding to fluorescein-labeled polyCAA RNA. The eIF3a and eIF3_R_ datasets are reproduced from Figure 3 for comparison. The anisotropy response observed for eIF3ac differs from that of isolated eIF3a and more closely resembles the response observed for eIF3_R_, indicating that inclusion of eIF3c alters eIF3a-driven RNA binding. **(B)** Extended fluorescence-anisotropy titration of isolated eIF3c with fluorescein-labeled polyCAA RNA. The anisotropy response did not reach saturation at eIF3c concentrations of up to 5 µM, consistent with weak RNA binding by isolated eIF3c. Points represent the mean of three independent titrations, and error bars represent the standard deviation.

**Supplemental Figure 7.**
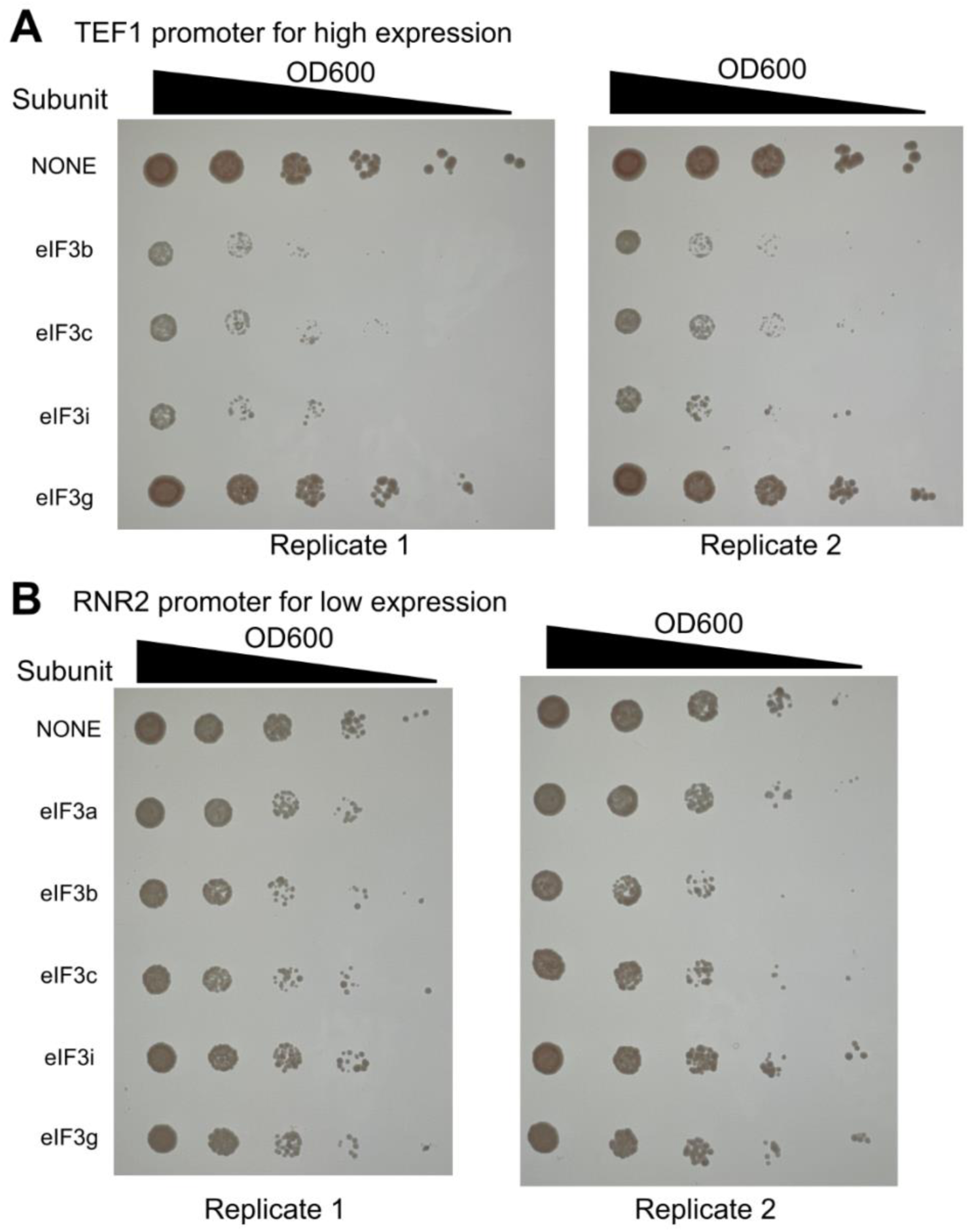
Plasmid-driven expression of individual eIF3 subunits alters yeast growth. **(A)** Spotting assays of *S. cerevisiae* cells carrying an empty vector control or plasmids expressing the indicated eIF3 subunits from the high-expression TEF1 promoter. Cultures were normalized to an OD_600_ of 0.5 and serially diluted 5-fold from left to right. Transformants carrying the TEF1-eIF3a construct could not be propagated and are therefore not shown. **(B)** Spotting assays of cells carrying an empty-vector control or plasmids expressing the indicated eIF3 subunits from the lower-expression RNR2 promoter. Two biological replicates are shown for each condition. Under these plasmid-expression conditions, growth defects were more pronounced with the TEF1 constructs than with the corresponding RNR2 constructs. The mechanisms underlying these phenotypes were not determined.

**Supplemental Figure 8.**
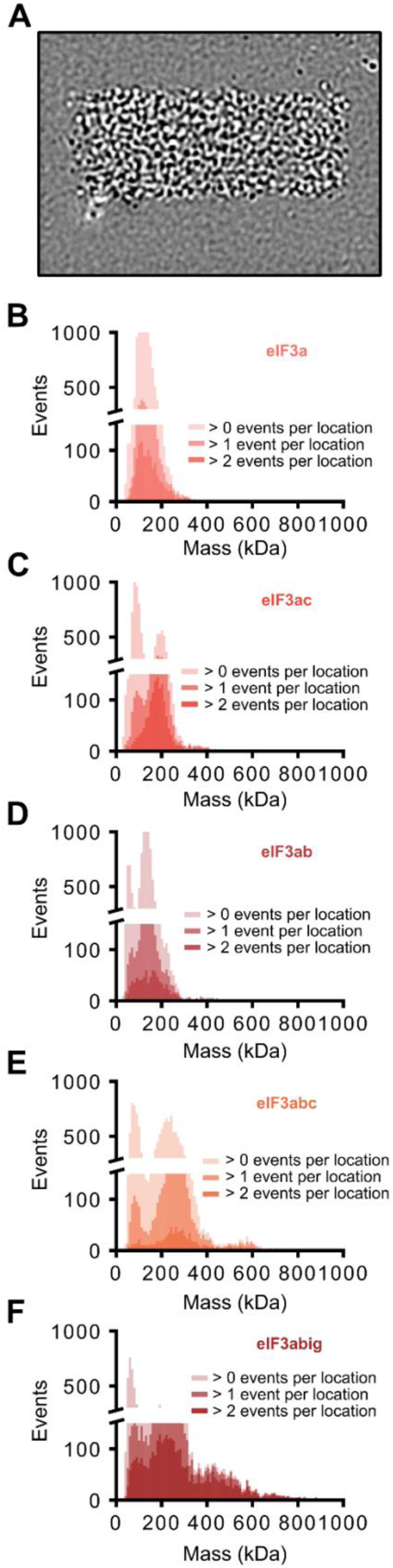
Repeated events observed for eIF3a and defined eIF3a-containing subcomplexes in the tethered-RNA mass-photometry assay. **(A)** Representative mass photometry frame acquired following laser-induced perturbation of the PEG-derivatized surface. **(B-F)** Apparent-mass distributions observed after addition of 200 nM **(B)** eIF3a, **(C)** eIF3ac, **(D)** eIF3ab, **(E)** eIF3abc, or **(F)** eIF3abig to PEG-derivatized coverslips bearing sparsely tethered biotinylated RNA. Distributions were filtered for locations exhibiting > 0, > 1, or > 2 events, as indicated. Repeated events were observed for eIF3a and each tested eIF3a-containing subcomplex, indicating that these species are competent to generate repeated events in the tethered-RNA assay. These measurements were not used to infer solution-phase composition, subunit-exchange behavior, molecular identities from apparent mass alone, or relative binding affinities.

**Supplemental Table 1.** Apparent equilibrium dissociation constants for RNA binding by eIF3 species. . Values are the mean *K*_d_^app^ ± standard deviation from three independent titrations and were obtained by fitting the fluorescence-anisotropy data to the quadratic binding equation described in the Methods. RNA lengths and sequences are provided in the Methods. *K*_d_^app^ values are reported only when the titration approached sufficient saturation to constrain the fit. N.B., no appreciable binding detected over the tested protein-concentration range; N.S., anisotropy response observed but insufficient saturation to determine a reliable K_d_^app^; **—**, experiment not performed.

| Apparent equilibrium dissociation constants, $K_d^{\text{app}}$ (nM) | | | | | |
| --- | --- | --- | --- | --- | --- |
|  | polyCAA (50nt) | polyCAA (45 nt) | polyCAA (40 nt) | polyUC (41 nt) | rpl41a-derived (51 nt) |
| eIF3<sub>N</sub> | 4.1 ± 1.3 | 8.3 ± 2.7 | N.B. | 3.3 ± 1.4 | 4.2 ± 0.2 |
| eIF3<sub>R</sub> | 47 ± 18 | 5.8 ± 0.7 | N.B. | 31 ± 12 | 101 ± 40 |
| eIF3a | 116 ± 55 | 45 ± 4.2 | N.B. | 73 ± 20 | 102 ± 28 |
| eIF3b | N.B. | — | — | N.B. | N.B. |
| eIF3c | N.S. | — | — | N.S. | N.S. |
| eIF3ig | N.B. | — | — | N.B. | N.B. |

$K_d^{\text{app}}$ assuming two-component, single-site binding N.B. no detectable binding over the concentration range tested N.S. nonsaturating response from which $K_d^{\text{app}}$ could not be reliably determined — Experiment not performed

